# Long-distance dispersal shaped the diversity of tribe Dorstenieae (Moraceae)

**DOI:** 10.1101/531855

**Authors:** Qian Zhang, Elliot Gardner, Nyree Zerega, Hervé Sauquet

## Abstract

**Aim:** The Neotropics have the highest terrestrial biodiversity on earth. Investigating the relationships between the floras of the Neotropics and other tropical areas is critical to understanding the origin and evolution of this mega-diverse region. Tribe Dorstenieae (Moraceae) has a pantropical distribution and almost equal number of species on both sides of the Atlantic. In this study, we investigate the relationship between the African and Neotropical floras using Dorstenieae (15 genera, 156 species, Moraceae) as a model clade.

**Location:** the Neotropics and Africa.

**Methods:** We used a targeted enrichment strategy with herbarium samples and a nuclear bait set to assemble a data set of 102 genes sampled from 83 (53%) species and fifteen genera (100%) of Dorstenieae, and five outgroup species. Phylogenetic relationships were reconstructed with maximum likelihood and coalescent approaches. This phylogeny was dated with a Bayesian relaxed clock model and four fossil calibrations. The biogeographic history of the group was then reconstructed with several dispersal-extinction-cladogenesis models (incl. DEC and DEC+J).

**Results:** The crown-group ages of Dorstenieae and *Dorstenia* were estimated in the Cretaceous (65.8-79.8 Ma) and the Paleocene (50.8-67.3 Ma), respectively. Tribe Dorstenieae as a whole appears to have originated in the joint area of continental Africa, Madagascar and Asia-Oceania area. The Neotropical species of *Dorstenia* diversified in the Eocene (29.8-44.7 Ma) and formed a clade nested within the African lineages in the genus. *Brosimum* s.l., with a crown-group age at the period of the Oligocene and Miocene (14.9-31.1 Ma), represents another Neotropical clade in Dorstenieae.

**Main conclusions:** Tribe Dorstenieae originated in the joint area of continental Africa, Madagascar and Asia-Oceania area in the Cretaceous and then dispersed into Neotropics twice. Neotropical diversification after long-distance dispersal across the Atlantic is the most plausible explanation for the extant distribution pattern of Dorstenieae.

## INTRODUCTION

The Neotropical ecozone has been defined as the region from central Mexico to southern Brazil (Morrone, 2014). The Neotropics hold the highest terrestrial biodiversity on earth (Antonelli & Sanmartín, 2011) and harbor all major tropical biomes: lowland rain forests, seasonally dry forests, mid-elevation montane forests, savannas, high elevation grasslands and deserts (Hughes et al., 2012). A recent estimate of tree species based on a pantropical tree inventory database suggested that the number of tree species in the Neotropics was as many as in the Indo-Pacific region and almost triple the counterparts in continental Africa (Slik et al., 2015). Several hypotheses have been proposed for the origins and evolution of Neotropical biodiversity during the past two decades. These hypotheses can be coarsely classified as biotic (e.g., dispersal ability, niche conservatism) and abiotic (time, climate, mountain uplift, Antonelli & Sanmartín, 2011).

After completely splitting from continental Africa in the Cretaceous (ca. 105 Ma) (McLoughlin, 2001), South America was isolated until the uplift of the Panama Isthmus (ca. 15 Ma), which connected North and South America (Montes et al., 2012; Bacon et al., 2015). Evidence of long-distance dispersals (LDD) among all of these three continents has been found with the development of molecular dating and phylogenetic approaches (Christenhusz & Chase, 2013). The relationships between the floras of the Neotropics and of other continents has intrigued researchers, as investigating these relationships not only can shed light on the origin and evolution of Neotropical biodiversity, but also help to understand disjunct distributions and long-distance dispersal. Taxonomically and genetically densely sampled phylogenetic analyses represent an ideal approach to improve the understanding of the origin and evolution of Neotropical diversity (Antonelli & Sanmartín, 2011; Hughes et al., 2012).

The generic composition of the angiosperm tribe Dorstenieae in the Moraceae family has been under recent scrutiny. Based on taxonomic treatments and recent molecular phylogenetic studies, Dorstenieae are currently thought to consist of fifteen genera (Table 1) and approximately 156 species (Berg et al. 2001; Clement & Weiblen 2009; Zerega et al. 2010; Chung et al. 2017). The most diverse genus in Dorstenieae is *Dorstenia*, which includes 113 species (Berg & Hijman, 1999; Mccoy & Massara, 2008; dos Santos & Neto, 2012; Machado & Marcelo Filho, 2012; Chase et al., 2013; dos Santos et al., 2013; Leal, 2014; Machado et al., 2014; Rzepecky, 2016). The genera in Dorstenieae are restricted to either side of the Atlantic, except *Dorstenia*, which has almost the same number of species in South America and continental Africa (50 in the Neotropics; 62 in continental Africa, Madagascar and Arabian Peninsula; 1 in India and Sri Lanka) (Berg & Hijman, 1999). In a recent phylogenetic study of Moraceae (Zhang et al., 2018), the relationships among the genera of Dorstenieae (except *Bosqueiopsis* and *Scyphosyce*) were reconstructed. Most of them were strongly supported, but the relationships of *Trilepisium*, *Brosimum* and *Treculia* remained unclear. The most densely sampled phylogenetic study of *Dorstenia* to date sampled 32 species (28%) and found *Dorstenia* to have originated in Africa, with three African species nested inside the Neotropical clade (Misiewicz & Zerega, 2012).

**Table 1.** Classification of Dorstenieae and the number of species for each genus.

| This study | No. of species<br>sampled in this<br>study /No. of<br>species | Distribution | Reference |
| --- | --- | --- | --- |
| *Allaeanthus* | 3/4 | D,E,F | (Chung et al., 2017) |
| *Bleekrodea* | 1/3 | D, F | (Rohwer & Berg, 1993) |
| *Bosqueiopsis* | 1/1 | C | (Rohwer & Berg, 1993;<br>Berg, 2001) |
| *Broussonetia* | 1/4 | F | (Rohwer & Berg, 1993;<br>Chung et al., 2017) |
| *Brosimum* | 10/15 | A, B | (Berg, 2001) |
| *Dorstenia* | 73/113 | A, B, C, D, E | (Berg & Hijman, 1999;<br>Berg, 2001; McCoy &<br>Massara, 2008; dos Santos<br>& Neto, 2012; Machado &<br>Marcelo Filho, 2012;<br>Chase et al., 2013; dos<br>Santos et al., 2013; Leal,<br>2014; Machado et al.,<br>2014; Rzepecky, 2016) |
| *Fatoua* | 1/2 | D, F | (Rohwer & Berg, 1993) |
| *Helianthostylis* | 1/2 | A | (Rohwer & Berg, 1993;<br>Berg, 2001) |
| *Malaisia* | 1/1 | F | (Wu et al., 2003; Clement<br>& Weiblen, 2009) |
| *Scyphosyce* | 2/2 | C | (Berg, 1977; Rohwer &<br>Berg, 1993) |
| *Sloetia* | 1/1 | F | (Clement & Weiblen,<br>2009; Tandang et al.,<br>2017) |
| *Treculia* | 3/3 | C, D | (Rohwer & Berg, 1993;<br>Zerega et al., 2010) |
| *Trilepisium* | 1/1 | C, D | (Rohwer & Berg, 1993;<br>Berg, 2001) |
| *Trymatococcus* | 1/2 | A | (Berg, 2001) |
| *Utsetela* | 1/2 | C | (Berg, 1977; Jongkind,<br>1995) |

Estimating absolute divergence times as accurately as possible is essential to biogeographic reconstruction to connect the evolutionary history of target taxa with past climate change and geographic events (Sauquet, 2013). Previous studies have estimated the crown-group age of Dorstenieae to be at least 71 Ma (Zerega et al., 2005), 50.6-72.5 Ma (Gardner et al., 2017), or 51.4-70.2 Ma (Zhang et al., 2018). While these estimates overlap, the crown-group age of *Dorstenia* has remained unclear. The crown-group age of *Dorstenia* has been estimated using a range of taxonomic sampling density to be 3.5-18.4 Ma (Zerega et al., 2005, with two Neotropical species included), 12.7-31.7 Ma (Zhang et al., 2018, with two neotropical and one African species included), or 84.8-132.0 Ma (Misiewicz & Zerega, 2012, with 15 neotropical and 14 African species included).

Approximately 10% of the species in Moraceae are herbaceous and all of them belong to Dorstenieae (more specifically all in *Dorstenia* and *Fatoua*; Berg, 2001). Species of Dorstenieae show diversity in pollination modes, dispersal mechanisms, and habit (Berg, 2001). The species of *Dorstenia* are found in a wide variety of habitats (e.g., tropical rain forest, savannas, or crevices of cliffs) and life forms (e.g., tree, shrubs, caulescent, herbaceous; Berg & Hijman, 1999; Berg, 2001; Misiewicz & Zerega, 2012). The pantropical distribution, almost equal diversity on both sides of the Atlantic, and the diverse traits of Dorstenieae make it a good model for understanding the origin and evolution of Neotropical biodiversity and the relationships between African and Neotropical floras.

Genomic targeted enrichment approaches have been shown to be more efficient and economic than Sanger sequencing (Lemmon & Lemmon, 2013; McKain et al., 2018) and have been widely used in phylogenetic studies in recent years (Xi et al., 2014; Fisher et al., 2016; Hart et al., 2016; Mitchell et al., 2017; Couvreur et al., 2018). In this study, we targeted nuclear genes as they have been suggested to hold the greatest potential for investigating the evolutionary history of angiosperms for several reasons. Firstly, nuclear genes have worked well in reconstructing more strongly supported phylogenetic relationships than organellar markers in both deep and shallow time scales (Xi et al., 2014; Mitchell et al., 2017). Secondly, nuclear genes are assumed to be unlinked, decreasing the probability of misleading phylogeny by part of the genes used (Fisher et al., 2016). Lastly, nuclear genes present multiple lineage histories, contrary to plastid genes, which are usually considered to represent a single locus.

In this study, we included samples representing all fifteen genera of Dorstenieae (Table 1) to reconstruct a comprehensive dated phylogenetic tree of Dorstenieae with two primary goals: 1) to test phylogenetic relationships and monophyly of Dorstenieae genera, and relationships within *Dorstenia*; and 2) to investigate divergence times and the biogeographic history of tribe Dorstenieae and genus *Dorstenia*. This study also provided an opportunity to test the potential of the targeted enrichment strategy to resolve species-level phylogenetic relationships using herbarium material, taking advantage of a recently developed nuclear bait set for Moraceae (Gardner et al., 2016, Johnson et al., 2016).

## MATERIALS AND METHODS

### Specimen and sample collection

In this study, we follow the classification of Clement & Weiblen (2009) with modifications based on Zerega et al. (2010, recommending *Treculia* be transferred to Dorstenieae) and Chung et al. (2017, reinstating *Allaeanthus* in Dorstenieae). This approach recognizes fifteen genera and approximately 153 species in this tribe (Table 1). We included 83 species (93 taxa) representing all currently recognized Dorstenieae genera and 54% of the species within tribe Dorstenieae (Table S1, we also collected samples of 22 more species in Dorstenieae and they were finally excluded in the main analyses, see below). Additionally, five outgroup taxa in Moraceae were included. They represent five of the six other Moraceae tribes: Artocarpeae, Castilleae, Ficeae, Moreae, and the newly created Parartocarpeae (Zerega & Gardner, 2019). Most taxa were extracted from herbarium material sampled from the Field Museum (F), the Missouri Botanical Garden (MO), the New York Botanical Garden (NY), and the Muséum national d’Histoire naturelle (P) (Table S1). Specimens collected within the last twenty years, and reliably identified material with inflorescences or infructescences were preferred. Samples include 55 (49%) species of *Dorstenia*, representing eight out of the nine sections proposed by Berg and Hijman (1999). Section *Bazzemia*, which contains only one species from Mozambique (Berg & Hijman, 1999), was not sampled.

### DNA extraction and sequencing

Whole genome DNA was extracted using a modified CTAB method (Doyle & Doyle, 1987). Extracted DNA was re-suspended in 50 μl light TE. The DNA concentration of each sample were measured using Qubit 2.0 fluorometer (Life Technologies, Carlsbad, CA, USA) following the standard protocol. For library preparation, we used 200 ng DNA when possible, but never less than 20 ng. All samples were run on an agarose gel to assess fragment size. Samples with DNA fragment lengths longer than 500 base pairs (bp) were sonicated on a Covaris M220 (Covaris, Wobum, MA, USA) for 45 seconds at 50 W peak power and a duty factor of 20%, which typically produces average fragment sizes of 550 bp. We prepared dual-indexed sequencing libraries using the KAPA Hyper Prep Library Construction Protocol (KAPA biosystems, Wilmington, MA, USA), generally following the manufacturer’s protocol except that most steps were performed at one-quarter volume to save costs. Low-input or very degraded samples were not size selected. Libraries were amplified using 12 cycles of PCR, but half of the unamplified template was retained in case PCR needed to be repeated. Products were cleaned using Solid Phase Reversible Immobilization (SPRI) and quantified using a high-sensitivity dsDNA assay on a Qubit 2.0. For samples with a concentration less than 5 μg/ml, we repeated PCR amplification with 14 cycles. Successful libraries were combined into seven pools of 13 to 14 libraries each. We hybridized the libraries to custom Moraceae probes (Gardner et al. 2016) manufacturered by Arbor Biosciences (Ann Arbor, MI, USA) as a MYbaits kit. Hybridization for 20 hours followed the manufacturer’s protocol, and products were reamplified 14 cycles of PCR. Amplified products were quantified on a Qubit 2.0, and fragment sizes were determined using a High-Sensitivity DNA assay on a BioAnalyzer 2100 (Agilent Technologies, Palo Alto, Califonia, USA). When required, adapter dimer was removed using 0.7x SPRI beads. All libraries except were then sequenced on a single lane of an Illumina HiSeq 2000 (2 × 100 bp, paired-end) by Genewiz (Genewiz, South Plainfield, NJ, USA).

To the 96 libraries prepared for this study, we added sequences from 14 samples prepared for other projects, including *Brosimum* (10)*, Trymatoccocus oligandrus*, *Helianthostylis sprucei*, *Allaeanthus luzonicus*, and *Malaisia scandens*, as well as five samples (*Artocarpus heterophyllus, Milicia excelsa, Parartocarpus venenosus, Ficus macrophylla*, and *Antiaropsis decipiens*) from Johnson et al. (2016) and Zerega and Gardner (Zerega & Gardner, 2019). These 20 samples were all sequenced on an Illumina MiSeq with somewhat longer reads (2×300bp, v3). Finally, we used transcriptomic reads for *Broussonetia papyrifera* obtained from GenBank.

### Sequence cleaning, assembly, and filtering

Sequences were assembled using HybPiper (Johnson et al., 2016), which uses reference sequences to guide local *de novo* assemblies of each target gene. Because Dorstenieae are phylogenetically distant from the baits sequences, which come from Artocarpeae and Moreae (Gardner et al. 2016), a new HybPiper reference was generated using six samples sequenced here, each with at least eight million read pairs (*Dorstenia bahiensis*, *D. cayapia*, *D. erythrandra*, *D. kameruniana*, *Treculia africana*, and *Fatoua villosa*). Reads were trimmed using Trimmomatic (Bolger et al., 2014) (LEADING:20 TRAILING:20 SLIDINGWINDOW:4:20 MINLEN:20) and assembled with SPAdes 3.10.1 (Bankevich et al., 2012) using default parameters. Coding sequences were predicted with Augustus (Stanke et al., 2004), using *Arabidopsis thaliana* genes as a reference. A seven-way orthology search was carried out with ProteinOrtho5 (Lechner et al., 2011) using all CDS over 200 bp from the *de novo* assemblies in addition to the *Artocarpus* HybPiper (Johnson et al., 2016) reference from Kates et al. (Kates et al., 2018). Orthologs present in at least three taxa were included in a new seven-taxon HybPiper reference consisting of in-frame CDS. This expanded reference contained approximately 500 genes.

Assembly of all reads then proceeded as follows. We first trimmed low quality bases and adapter sequences using Trimmomatic (Bolger et al., 2014) with the following parameters: ILLUMINACLIP:TruSeq3-PE-2.fa:2:30:10 LEADING:20 TRAILING:20 SLIDINGWINDOW:4:20 MINLEN:40. To ensure that quality trimming worked as expected, we examined a subset of reads were both before and after trimming with FastQC (https://www.bioinformatics.babraham.ac.uk/projects/fastqc/). Reference-guided assembly then proceeded with HybPiper (Johnson et al., 2016), using the new reference described above. Briefly, the program works as follows: reads are sorted by gene based on the reference. Local *de novo* assemblies are carried out using SPAdes (Bankevich et al., 2012), and coding DNA sequences (CDS) are predicted using Exonerate (Slater & Birney, 2005). When a gene is assembled into several disconnected contigs—common in degraded samples where the fragments (and therefore effective read length) are very short—HybPiper scaffolds these short contigs in the correct order based on the reference. In the event that multiple genes are assembled for a single target, HybPiper distinguishes orthologs from paralogs using a combination of alignment length and identity relative to the reference. HybPiper outputs include in-frame CDS sequences as well as “supercontigs,” which contain CDS as well as any flanking non-coding sequences. For our analyses, we used only the supercontig sequences. We then filtered the sequences within loci to remove those less than 150 bp long or shorter 25% of the average length for the locus.

After filtering, the number of genes recovered for each taxon varied widely among taxa (from zero for *D. scaphigera* and 515 for *Ficus macrophylla*) and the number of taxa retrieved for each gene differed sharply among genes (from two to 101). These extreme differences would result in data set with a high proportion of missing data, which could impact the accuracy of both phylogenetic reconstruction and divergence time estimate. Therefore, we filtered the data set further by selecting 102 genes with high taxon occupancy and excluding taxa for which less than 30 of these genes were recovered (Table S1).

### Phylogenomic reconstruction and molecular dating

Sequences were aligned with MACSE (Ranwez et al., 2011), gene by gene (-fs 30 -stop 50). We reconstructed phylogenetic relationships using both a concatenated maximum likelihood (ML) and a coalescent approach. *Milicia exceisa* and *Artocarpus venenosus* were specified as the most external outgroups based on recent phylogenetic analyses of Moraceae (Clement & Weiblen, 2009; Zerega et al., 2010; Chung et al., 2017; Gardner et al., 2017; Zhang et al., 2018). Sequences were concatenated into a supermatrix and partitioned according to genes using the fasta_merge.py script from HybPiper; a ML tree was reconstructed with RAxML v8.2.10 (Stamatakis, 2014) as implemented on CIPRES (Miller et al., 2010) 1000 rapid bootstrap replicates. GTRGAMMA was chosen as the substitution model. For the coalescent approach, we first reconstructed ML trees with RAxML v8.2.10 for each gene, using with GTRGAMMA model and 200 rapid bootstrap replicates. These ML trees for each gene were used to infer a species tree using the summary coalescent approach implemented in ASTRAL-II (Mirarab & Warnow, 2015). Node support was calculated based on 200 bootstrap replicates, with resampling within loci.

Because the topologies of the ML and species trees were broadly consistent, we used the ML tree to estimate divergence times with a relaxed clock model and four fossil calibrations from Zhang et al. (2018). The root age (crown-group node of Moraceae) was constrained between 73 to 85 Ma, based on the results of Zhang et al. (2018) who used a more comprehensive sample of species within and outside Moraceae along with 12 fossil age constraints. The fossil wood of *Artocarpoxylon deccanensis* Mehrotra, Prakash, and Bande (at least 64.0 Ma) (Mehrotra et al., 1984) was used to calibrate the split between *Artocarpus* and *Milicia* as we used *Artocarpus heterophyllus* and *Milicia excelsa* to represent Artocarpeae and Moreae, respectively. The fossil fruits *Morus tymensis* Dorofeev (at least 33.9 Ma) (Collinson, 1989) used to calibrate Moreae in Zhang et al. (2018) would here provide an additional, but younger (and hence uninformative) minimum age constraint on the stem node of *Milicia*. The fossil endocarps of *Broussonetia rugosa* Chandler (Chandler, 1961) were used to constrain the crown group of *Broussonetia* s.l. (incl. *Allaeanthus*, previously included in *Broussonetia*, and *Malaisia*; Chung et al., 2017) to a minimum age of 33.9 Ma. The fossil achenes of *Ficus* (*F. lucidus* Chandler) (Chandler, 1962) were used to calibrate the stem group of *Ficus* with a minimum age of 56.0 Ma. Thus, we used one secondary calibration (root) and three fossil age constraints in our analyses.

Both penalized likelihood (PL) and Bayesian relaxed clock approaches were used to estimate divergence times in Dorstenieae. PL was implemented in r8s v1.7 (Sanderson, 2003) with strict minimum and maximum age constraints as described above. For the PL approach, the best smoothing value was first determined using cross validation by testing 21 values of smoothing parameter scaling from 0.1 to 1000. The optimum value (i.e., with lowest chi-square) of 1.6 was then used as the smoothing parameter in divergence time estimation.

For the Bayesian approach, we used MCMCTree as implemented in the PAML v4.9 package (Yang, 2007). MCMCTree has been used to estimate divergence times with other similar phylogenomic datasets (e.g., Foster et al., 2017), where other programs such as BEAST2 (Bouckaert et al., 2014) would take too long to converge. The tree topology was fixed, using the best-scoring tree from the RAxML analysis, and all calibrations were implemented as uniform priors with soft boundaries (2.5% on both sides) (Ho & Phillips, 2009). We set the minimum age of the fossil and as the minimum boundary and the younger bound of the root (73 Ma) as the maximum boundary of the uniform distribution for each fossil calibration. The birth-death process was used as the tree prior. For the MCMCTree analysis, we used the supermatrix, analyzed as a single partition under the GTR substitution model.

We first ran baseml in PAML v4.9 with a strict clock model to estimate the rough mean of parameters such as the shape parameter for the overall rate and the transition/transversion rate ratio. Two steps are needed for divergence time estimation by approximate likelihood in MCMCTree. We first estimated the gradient and Hessian, and then used them to estimate the divergence times. We set the prior in both steps according to the estimates of baseml. In both steps, we ran the process for 38.5 million generations, with the first 10% of the chain length discarded as burnin, sampling a total of 10,000 generations at a frequency of once very 3,500 generations. Two independent runs with the same settings were conducted to confirm the convergence of the MCMC. To check the influence of the prior on the estimation, we used another prior setting (program defaults), followed the same steps as above, then ran the chain for 22 million generations, sampling a total of 10,000 generations at a frequency of once every 2,000 generations. After checking the convergence of the runs, we combined the results of them for each prior setting. Two additional independent runs with the two different prior settings, but without data, were also conducted to test the impact of the priors on the results. The convergence of Markov chain Monte Carlo (MCMC) was checked by reading the result log files in Tracer v1.7 (Rambaut et al., 2018). For the Bayesian approach, divergence was checked by confirming that the effective sample size (ESS) of parameters in the two independent runs was over 100 after removing the first 10% of chain length as burnin.

### Reconstruction of biogeographic history

Distribution data for the taxa sampled were collected from monographs and revisions (Rohwer & Berg, 1993; Berg & Hijman, 1999; Berg, 2001; Chung et al., 2017). We separated the distribution area of extant species of Dorstenieae into six areas (Figure 1) following a recent study of Annonaceae, which share a similar global distribution (Couvreur et al., 2011): A, South America; B, North/Central America; C, Africa; D, Madagascar; E, India and Sri Lanka; F, Southeast Asia and Oceania. We decided to combine the two areas west and east of Wallace’s line (areas F and G of Couvreur et al., 2011) into a single area (area F) because all species of Dorstenieae distributed east of Wallace’s line were also observed west of Wallace’s line in our data set (e.g., *Allaeanthus luzonicus*, *Fatoua villosa*).

**Figure 1.**
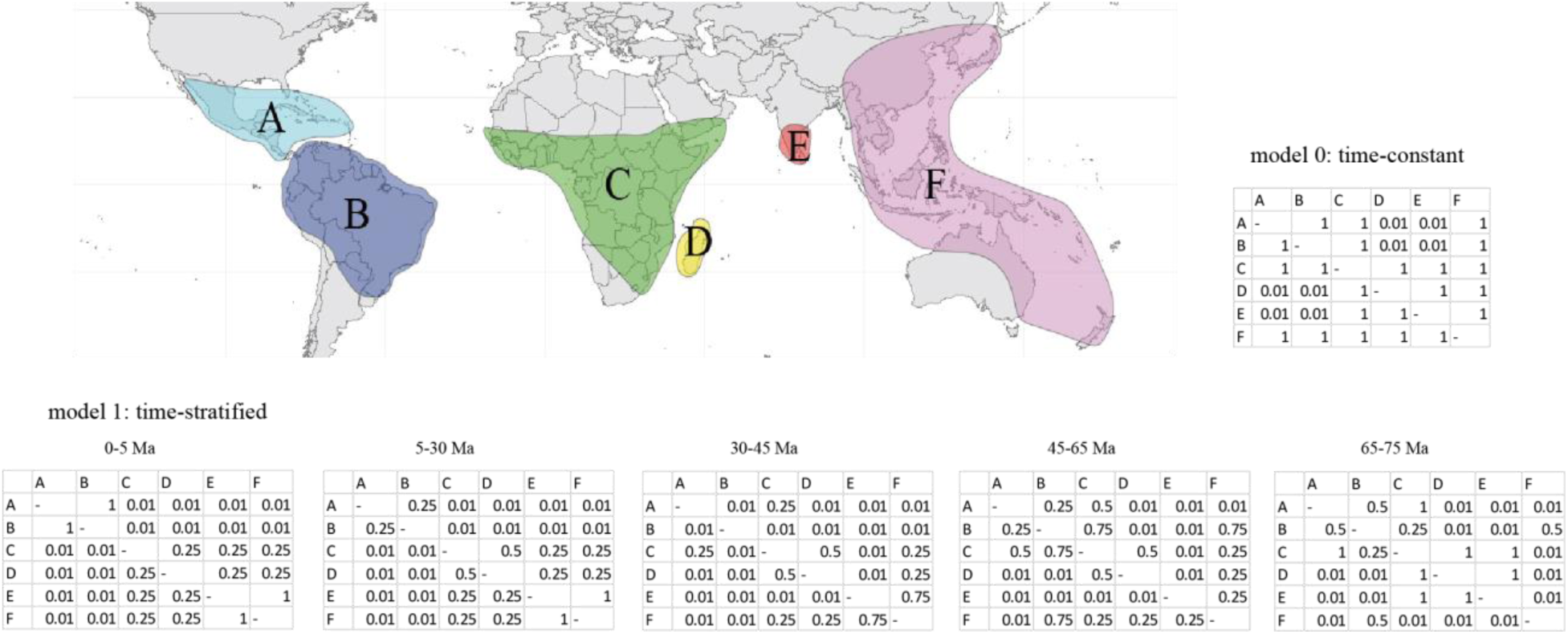
Delimitation of distribution areas of Dorstenieae and relative dispersal matrices for the time-constant (model 0) and time-stratified (model 1) models. A, South America; B, North/Central America; C, Africa; D, Madagascar; E, India and Sri Lanka; F, Southeast Asia, and Oceania. Five levels of dispersal probability, 0.01, 0.25, 0.5, 0.75 and 1, representing the probability from low to high.

Only Dorstenieae (93 taxa, 83 species) were kept in the biogeographic reconstruction to avoid the bias of incompletely sampled outgroups. BioGeoBEARS (Matzke, 2013a) in R v3.5.1 (R Core Team, 2018) was used to reconstruct the biogeographic history of the clade with the DEC and DEC+J models. Prior studies have suggested that the DEC+J model can exacerbate the bias of preferring cladogenetic events (i.e., sympatry, vicariance and founder event speciation) over anagenetic processes in the DEC model, implying that the higher likelihoods typically obtained with DEC+J do not necessarily mean a better fit to the data than DEC (Ree & Sanmartín, 2018). Therefore, we employed both the DEC and the DEC+J models. In addition, we ran the analyses with the DIVA-like and BAYAREA-like models including or not including founder-event speciation (+J). All analyses were run both with a simple dispersal matrix and a time-stratified model, hereafter referred to as model 0 and model 1, respectively. Both models were inspired by Couvreur et al. (2011) and are constrained by unequal dispersal relative rates aimed at reflecting the connectivity of biogeographic areas (Figure 1). In the time-stratified model, the constraints on dispersal probabilities vary through five time periods (0-5, 5-30, 30-45, 45-65, and 65-75 Ma) according to physical distance between areas (Figure 1). For instance, the dispersal rates from continental Africa (C) and Madagascar (D) to India and Sri Lanka (E) were constrained to be very low during the rafting of the Indian plate 30-65 Ma. Because all sampled species of Dorstenieae currently occupy no more than two areas, the maximum number of co-occurrence areas was set as three in all analyses.

## RESULTS

### Sequencing

Sequencing and assembly statistics appear in Table S1. Enrichment ranged from 0.34% to 84% reads on target, with the enrichment found in Artocarpeae and Moreae (58% and 84% on target, respectively) and the least efficient found in the genus *Dorstenia* (0.34% on target). The final data set consisted of 98 taxa (89 species, spanning all of the currently recognized genera) and 102 genes. The width of the aligned supermatrix was 132,753 bp with 29.51% gaps and missing data.

### Phylogenetic relationships

The maximum likelihood (ML) and coalescent analyses produced broadly identical topologies, with some weakly supported differences at shallow phylogenetic depths (Figure 2, Figure S1a,b). Most of the nodes on the tree were strongly supported (over 90% bootstrap support, Figure 2). *Fatoua villosa* was found to be sister to the remaining of Dorstenieae. *Malaisia scandens* and *Broussonetia papyrifera* together formed a clade that is sister to *Allaeanthus*. All genera of Dorstenieae sampled more than once were found to be monophyletic with two notable exceptions. *Trymatococcus oligandrus* and *Helianthostylis sprucei* appear to be nested in *Brosimum*, and *Scyphosyce* (two species) and *Utsetela gabonensis* were found to be nested among early-diverging lineages of *Dorstenia*. Hereafter we refer to the clade of *Brosimum*, *Trymatococcus* and *Helianthostylis* as *Brosimum* s.l., and to the clade of *Dorstenia*, *Scyphosyce* and *Utsetela* as *Dorstenia* s.l. The Neotropical species of *Dorstenia* formed a clade well nested among African lineages. *Dorstenia elliptica* (from Central Africa) appears to be sister to this Neotropical *Dorstenia* clade. The Central/North American species of *Dorstenia* formed two clades nested among the South American lineages with strong support as well. Internal branch lengths were comparatively shorter in Neotropical *Dorstenia* and most of the differences between the ML and coalescent approach concentrated in this clade. Short branches were also observed in *Brosimum* s.l., the other Neotropical clade in Dorstenieae, but in this case both reconstruction approaches showed identical topologies. Six species had duplicated samples in this study, four of them were sister or close to the other samples of the same species, while sample *D. brasiliensis*-1 and *D. arifolia* were not (but still lie in the Neotropical clade). Although in similar positions on the phylogenetic trees reconstructed by two approaches, the support for the splits of these two species were low in the coalescent tree (bootstrap support less than 50%) but high in the maximum likelihood tree (bootstrap support over 90%). Several nodes were comparatively weakly supported in *Dorstenia* (Figure 2). Our samples represented eight out of nine recognized sections in *Dorstenia*. None of them was found to be monophyletic in our analyses (Figure S1). All of the Neotropical species of Dorstenieae outside of *Dorstenia* also formed a clade (i.e., *Brosimum* s.l., incl. *Trymatococcus* and *Helianthostylis*).

**Figure 2.**
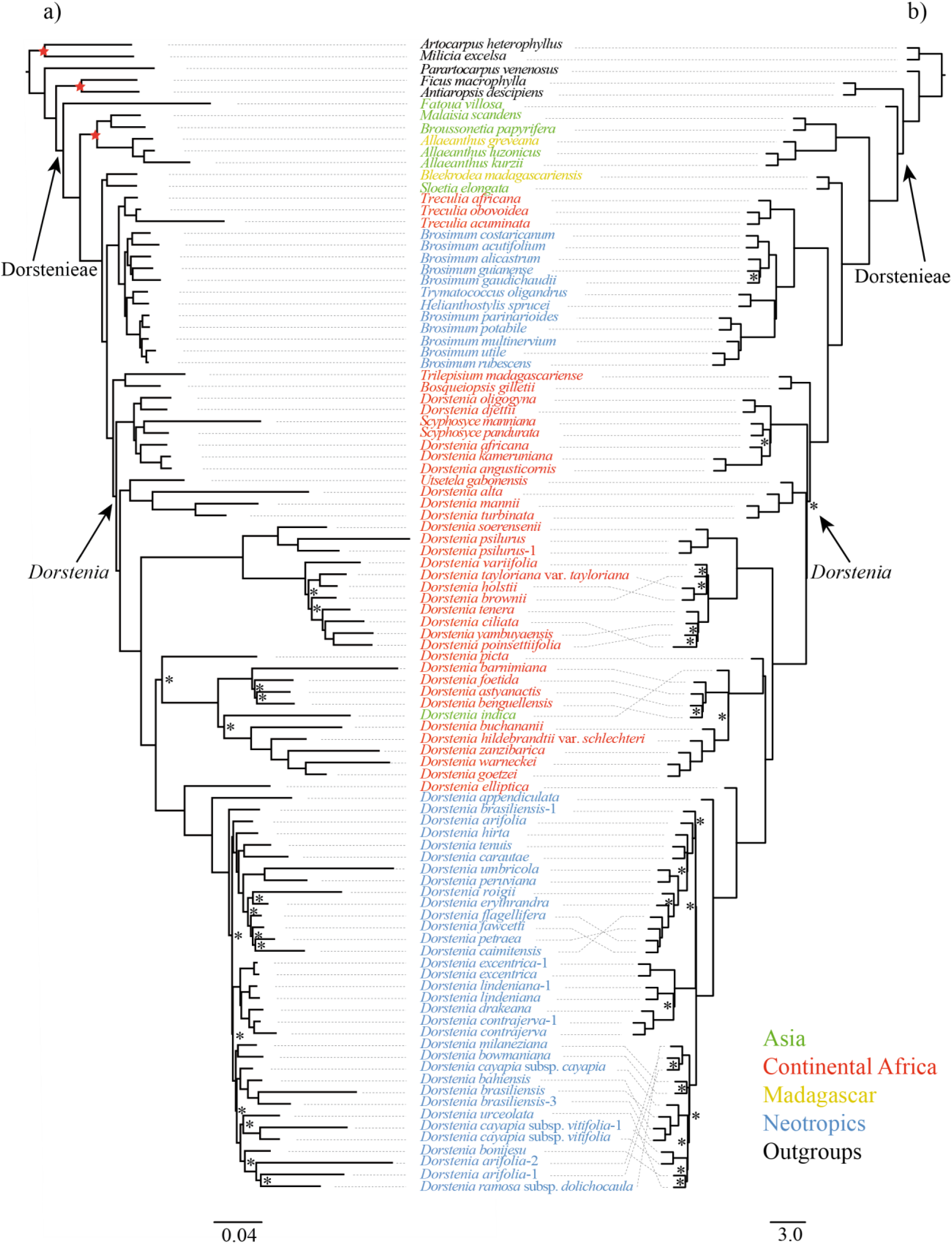
Maximum Likelihood phylogenomic tree (a) and ASTRAL tree (b) of Dorstenieae. Fossil-calibrated nodes are marked as red stars. Nodes with bootstrap support value (less than 90% are indicated with an asterisk. Tip names are colored by general distribution area.

### Divergence times of Dorstenieae and *Dorstenia*

The crown-group age of tribe Dorstenieae was estimated in the Cretaceous (65.8-79.8 Ma) and that of *Dorstenia* in the Paleocene (50.8-67.3 Ma) (Table 2, Figure S2). The stem and crown-group ages of the Neotropical *Dorstenia* clade were dated in the Eocene to early Oligocene (34.6-51.8 Ma and 29.8-44.7 Ma, respectively). *Brosimum* s.l., the other Neotropical clade in Dorstenieae, was estimated to date from the late Eocene to early Miocene (stem node: 19.6-41.5 Ma; crown node: 14.9-31.1 Ma). Runs with the prior only showed different results to those with data, indicating the data had a significant impact on the posterior (results not shown). Results of the two different prior settings were similar to each other (Table 2, Figure S2a,b). Divergence times estimated with the PL approach were all compatible with those from the Bayesian approach (i.e., falling within the 95% credibility intervals) except the age of crown-group *Broussonetia* s.l., which was significantly older with PL (Table 2, Figure S2c).

**Table 2.** Divergence time estimates with penalized likelihood (PL with r8s) and Bayesian (MCMCTree) approaches for key nodes of Dorstenieae.

|  | r8s (Ma) | MCMCTree set1 (Ma) | MCMCTree set2 (Ma) |
| --- | --- | --- | --- |
| SG Dorstenieae | 77.3 | 68.7-82.1 | 68.9-82.1 |
| CG Dorstenieae | 75.4 | 65.2-79.8 | 65.8-79.8 |
| CG *Treculia* | 35.3 | 10.9-32.7 | 11.2-32.4 |
| SG *Brosimum* s.l. | 40.7 | 19.4-42.9 | 19.6-41.5 |
| CG *Brosimum* s.l. | 27.9 | 14.7-31.7 | 14.9-31.1 |
| SG *Broussonetia* s.l. | 70.4 | 60.2-76.4 | 61.1-76.2 |
| CG *Broussonetia* s.l. | 52 | 30.5-41.8 | 30.2-40.7 |
| CG *Scyphosyce* | 34.7 | 10.8-31.5 | 10.7-30.9 |
| SG *Dorstenia* | 61.9 | 51.2-68.5 | 52.3-68.7 |
| CG *Dorstenia* | 61.3 | 49.8-67.0 | 50.8-67.3 |
| SG *Dorstenia* Neo | 43.4 | 34.3-52.1 | 34.6-51.8 |
| CG *Dorstenia* Neo | 33.4 | 29.7-44.7 | 29.8-44.7 |
| SG *Dorstenia* MAm clade1 | 23.1 | 16.5-28.2 | 16.7-28.0 |
| CG *Dorstenia* MAm clade1 | 20.2 | 12.6-24.9 | 12.8-24.6 |
| SG *Dorstenia* MAm clade2 | 25.4 | 23.9-35.1 | 24.1-34.7 |
| CG *Dorstenia* MAm clade2 | 13.3 | 12.1-28.9 | 12.0-28.4 |

CG: crown group; SG: stem group MCMCTree set1: results of estimate in MCMCTree with the prior setting referring to the results of baseml MCMCTree set2: results of estimate with the default prior setting in MCMCTree *Brosimum* s.l. includes *Brosimum*, *Trymatococus* and *Helianthostylis* *Broussonetia* s.l. includes *Broussonetia*, *Malaisia*, and *Allaeanthus* *Dorstenia* Neo: Neotropical *Dorstenia* species *Dorstenia* MAm Clade1: *Dorstenia* species distributed in Central and North America, from *D. erythrandra* to *D. caimitensis* in Figure 3 *Dorstenia* MAm Clade2: *Dorstenia* species distributed in Central and North America, from *D. excentrica* to *D. contrajerva* in Figure 3

### Biogeographic history of Dorstenieae

We analyzed the data set with both the dispersal-extinction-cladogenesis (DEC) and DEC+J (DEC with founder-event speciation) models. The time-stratified model fit the data significantly better than the time-constant model in all analyses conducted (Table S2), therefore we focus mainly on the results from the time-stratified analyses here, unless otherwise mentioned. In addition, the DEC+J model had a lower value of Akaike information criterion (AIC) than that of DEC (Table S2). The ancestral distribution area of Dorstenieae was estimated to be the combined area of continental Africa, Madagascar, and Southeast Asia and Oceania (CDF) with both the DEC and DEC+J models (Figure 3, Figure S3c). The ancestral area for both the stem and crown-group nodes of *Dorstenia* s.l. (incl. *Scyphosyce* and *Utsetela*) were estimated in continental Africa (C) with both models. South America (A) was found as the ancestral area of both the stem and crown-group nodes of the Neotropical *Dorstenia* clade with the DEC+J model (Figure 3), while the DEC model reconstructed the combined area of South America and continental Africa (AC) as ancestral for the stem-group node of this clade (Figure S3c). *Dorstenia indica*, endemic to India and Sri Lanka, was found to be nested in an African clade and diverged from its sister group in the Eocene to Oligocene (26.2-46.7 Ma). The Central/North American *Dorstenia* species clustered into two clades with strong support (Figure 2), suggesting two independent colonizations from South America to Central/North America during the Oligocene to Miocene (Table 2). The ancestral states of the crown-group nodes of both Central/North American clades were estimated as Central/North American. The stem-group nodes of clade 1 (from *D. erythrandra* to *D. caimitensis*) and clade 2 (from *D. excentrica* to *D. contrajerva*) were estimated to be Central/North American and South American, respectively (Figure 3). The stem and crown-group nodes of *Brosimum* s.l. were estimated in the joint area of South America and continental Africa (AC), and in South America (A), respectively, with both the DEC and DEC+J models.

**Figure 3.**
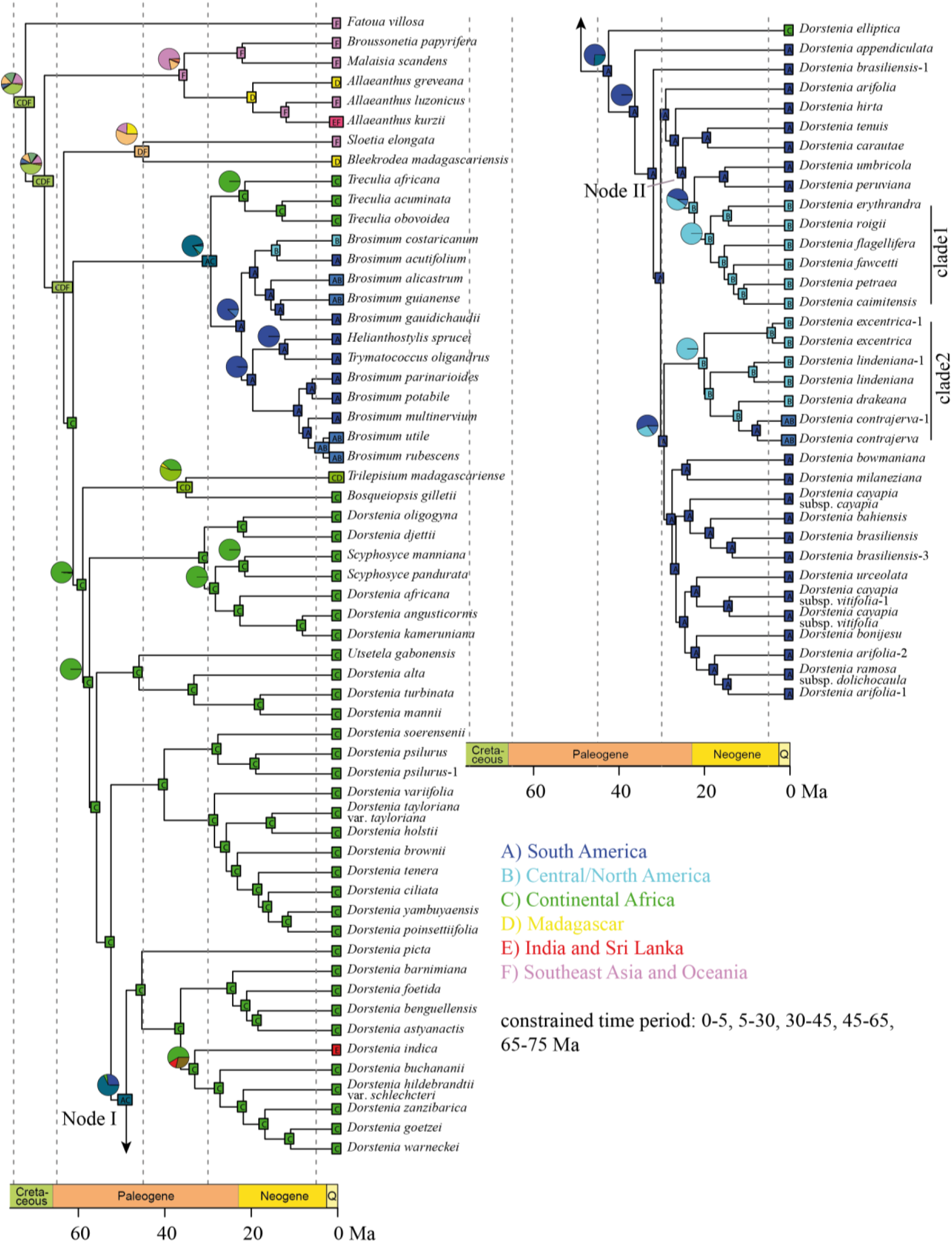
Biogeographic history reconstruction of Dorstenieae based on the time-stratified DEC+J model. Inferred ancestral distribution areas prior to speciation are indicated on the nodes. Pie charts for selected nodes represent the relative probability (proportional likelihoods) of alternative ancestral areas (for full details, see Figure S3d).

The additional biogeographic models produced similar results to those of the DEC and DEC+J models, with a few exceptions (Figure S3, Table S3). Reconstruction with time-constant models showed similar results as with time-stratified models. The ancestral distribution area for the crown-group node of Dorstenieae was estimated to be Southeast Asia and Oceania (F) by all the models based on the DIVA-like model (Figure S3i-l). In the time-stratified BayArea-like models, with or without founder-event speciation, the ancestral area of the crown-group node of Dorstenieae was estimated as the joint area of continental Africa and Southeast Asia and Oceania (CF), or as Southeast Asia and Oceania (F), respectively. The same pattern was found in reconstructions with time-constant BayArea-like models (Figure S3e-h). The stem and crown-group nodes of *Dorstenia* were estimated in continental Africa (C) in all the models. The crown-group node of Neotropical *Dorstenia* was estimated in South America (A) in all analyses, while the stem-group node was different among models. All the BayArea-like models estimated this node in continental Africa (C) (Figure S3e-h). The time-stratified DIVA-like models estimated the joint area of South America and continental Africa (AC), or South America (A) alone as the ancestral area of this node, with or without founder-event speciation respectively. The same results were found in time-constant DIVA-like models (Figure S3i-l). Lastly, we also ran all biogeographic analyses with the chronograms reconstructed with the penalized likelihood approach. The results were similar with some exceptions (results not shown). For instance, Southeast Asia and Oceania (F) was estimated as the ancestral area of the crown-group node of Dorstenieae with the time-stratified DEC+J model. The stem-group node of Neotropical *Dorstenia* was estimated as continental Africa (C) by the same model.

## DISCUSSION

### Success of the targeted enrichment strategy with herbarium specimens

Most (99%) of the samples in this study were from herbarium specimens (Table S1). Some of them were collected more than 40 years ago and the amount of sample collections from the herbarium was typically limited (around 3 to 20 mg) due to destructive sampling policies. Three samples (*Dorstenia aristeguietae*, *D. choconiana*, and *D. prorepens*) were filtered by HybPiper because of the low matching of reads to the reference. We excluded another fifteen taxa for the low number of genes recovered (less than 30, Table S1). The lowest amount of DNA used among the final 98 taxa retained in our dataset was 25.1 ng (*Dorstenia brasiliensis*). Samples that were excluded ranged from being over 100 years old to 11 years old, while samples that were included were collected as long ago as 1923. These results suggest that the degradation of DNA in old herbarium specimens appears to have had little influence on this study. This may be because of the short DNA fragments (on average less than 500 bp long) needed in the library preparation, those longer than this size requiring sonication. Thus, our results suggest that the targeted enrichment strategy and HybPiper pipeline (Johnson et al., 2016) worked well with a broad range of ages of herbarium specimens. Sequencing herbarium samples is a valuable approach for phylogenetics, as many species can be difficult to collect. Herbarium specimens often represent reliable and accurate vouchers of species identification, and some species may be rare or have even gone extinct in the wild (Särkinen et al., 2012; Staats et al., 2013). The success of similar targeted enrichment strategies with historical specimens for phylogenetic studies has previously been highlighted in other lineages of angiosperms at various scales, including *Arabidopsis thaliana* (Brassicaceae,Staats et al., 2013), *Inga* (Fabaceae, Hart et al., 2016), and Annonaceae.

Our results also suggest that the baits, which were originally designed for tribes Artocarpeae and Moreae in the same family (Gardner et al., 2016), worked well in Dorstenieae. In addition to the phylogenetic markers developed from 333 inferred single-copy exons for Moraceae, we retrieved approximately another 200 genes in this study for Dorstenieae. Those assembled untargeted genes had a mean identity to at least one target gene of 84%, and a mean alignment length to at least one target gene of 87% (as a percentage of the untargeted genes). These Moraceae specific baits worked well throughout the entire Moraceae family (Zerega & Gardner, 2019). Five untargeted genes were included in our final 102-gene data set. The most genes retrieved for one taxon was 515 from the outgroup sample *Ficus macrophylla*, suggesting a high probability that these baits would also work well to explore relationships within Ficeae (*Ficus*), the largest tribe in Moraceae (Couvreur et al., 2019).

### Phylogenetic relationships in Dorstenieae and *Dorstenia*

With all the extant genera of Dorstenieae included, this is the most densely sampled phylogenetic study of Dorstenieae to date. The reconstructed relationships (Figure 2) are generally consistent with previous work (Zerega et al., 2005; Clement & Weiblen, 2009; Misiewicz & Zerega, 2012; Chung et al., 2017; Zhang et al., 2018), but with stronger support, especially in *Brosimum* s.l. and *Dorstenia* s.l..

Prior to this study, the most densely sampled molecular phylogenetic analysis of *Dorstenia* was provided by Misiewicz and Zerega (2012), based on ITS sequences of 35 taxa (32 species) of *Dorstenia* and seven outgroup species. Our results are similar to those of Misiewicz and Zerega (2012) with respect to shallow-level relationships. However, they differ markedly at a deeper level in that the authors had found three African species (*D. variifolia*, *D. tayloriana* var. *tayloriana*, and *D. cuspidata*) to be nested in the Neotropical clade of *Dorstenia*. In our analyses, *D. variifolia* and *D. tayloriana* were also sampled and they were sister species as in Misiewicz & Zerega (2012), but we found these species nested in African clades with strong support. The difference may be explained by the root setting methods, the variation in each study of both number of genes and density of taxon sampling for both *Dorstenia* species as well as of non-*Dorstenia* species within the tribe Dorstenieae. Most differences between the ML and coalescent trees reconstructed in this study concentrated in the Neotropical *Dorstenia* clade (Figure 2). This result and the short branches observed in this clade suggest incomplete lineage sorting in the diversification of Neotropical *Dorstenia* species (Pamilo & Nei, 1988).

### Divergence times of Dorstenieae and its genera

Runs with or without data showed different results suggesting the data were informative. Runs with different prior settings showed similar estimates, suggesting that our results are robust to various assumptions on rate variation. The crown-group age of Dorstenieae was estimated in the Upper Cretaceous (65.8-79.8 Ma; Table 2), which overlaps with former studies that used fewer genes (Zerega et al., 2005; Gardner et al., 2017; Zhang et al., 2018). The crown-group age of *Dorstenia* was estimated in the Paleocene (50.8-67.3 Ma), which is younger than in Misiewicz and Zerega (2012). Although the estimated stem-group node of Neotropical *Dorstenia* clade in our study was younger than that in Misiewicz and Zerega (2012), the crown-group node of the same clade fell within a similar range in both studies. The difference in the stem-group node may be caused by different topologies and number of genes used to estimate the divergence time. The crown-group age of *Brosimum* s.l. and *Broussonetia* s.l. were estimated in the period from Oligocene to Miocene and the Eocene (14.9-31.1 Ma and 30.2-40.7 Ma, respectively), during which time the whole earth cooled down from Mid-Eocene Climatic Optimum and was warmer than the current climate (Zachos et al., 2008).

### Biogeographic history of Dorstenieae

Our results suggest that the most recent common ancestor of Dorstenieae was widely distributed in the joint area of continental Africa, Madagascar and Southeast Asia and Oceania (CDF, Figure 3, S3) in the Cretaceous (65.8-79.8 Ma), during which time these three areas were already separated from one another (PALEOMAP project, http://www.scotese.com/). Subsequently, at least two dispersals to South America (*Brosimum* s.l. and Neotropical *Dorstenia*) occurred during the evolutionary history of Dorstenieae. In our analyses, the stem-group ages of Dorstenieae (68.9-82.1 Ma), *Brosimum* s.l. (19.6-41.5 Ma) and Neotropical *Dorstenia* (34.6-51.8 Ma) were all estimated to be younger than the separation of South America and continental Africa at ca. 105 Ma (McLoughlin, 2001), suggesting that vicariance caused by Gondwanan breakup is unlikely to have played a role in the diversification of the two Neotropical clades. Long-distance dispersal, which is an indispensable process in Neotropical flora assembly (Hughes et al., 2012) may instead explain the origin of the two Neotropical clades in Dorstenieae.

The dispersal (range expansion) at the origin of the Neotropical clade of *Dorstenia* is inferred to have occurred from the Paleocene to early Eocene (time period between the stem and crown-group nodes of Node I, 42.1-62.1 Ma, Figure 3, S2, S3c), during which time angiosperms were already significantly more diverse in the Neotropics than in earlier time periods, according to palynological evidence (Jaramillo et al., 2006). During this period, global temperature increased, leading to the middle Eocene climatic optimum (Zachos et al., 2008). The increasing temperature has been suggested as one of the factors for the extension of the Neotropical region in the Eocene (Hughes et al., 2012). A larger Neotropical area at that time would have increased the probability of the successful colonization of species from Africa to a suitable habitat in the Neotropics. Curiously, in these reconstructions, *D. elliptica* (an African species sister to the Neotropical *Dorstenia* clade) was inferred to be the result of a back dispersal from South America to continental Africa (Figure 3, S3c). Whether this somewhat unexpected result is plausible or an example of pathological behavior of DEC models (Ree & Sanmartín, 2018) will require further investigation.

Two independent dispersals from South America to Central/North America in *Dorstenia* were estimated from the Oligocene to Miocene (period from Node II to stem-group node of Clade 1, 16.7-30.6 Ma and period from stem to crown group of Clade 2, 12.0-34.7 Ma), during which time waves of dispersal from South to North America have been found in other lineages (Bacon et al., 2015). This result supports the dispersal of plants from South America to Central/North America in the Neogene (Bagley & Johnson, 2014). The branch length or time between the stem and crown group of Neotropical *Dorstenia* was very short (less than 10 Ma, Table 2), suggesting the rapid divergence of the clade. The branch length between the stem and crown group of the other Neotropical Dorstenieae lineage (*Brosimum* s.l.) was also short (ca. 12 Ma, Table 2). Therefore, LDD followed by rapid diversification would explain the extant distribution pattern of Neotropical Dorstenieae species. A similar pathway was found in the pantropically distributed tribe Annoneae (Annonaceae) (Thomas et al., 2017; Williams et al., 2017).

A diversity of seed dispersal modes has been reported in Dorstenieae, including autochory by expulsion or ejection of endocarp in *Dorstenia*, *Bleekrodea*, *Fatoua*, and zoochory in *Brosimum lactescens* and *Trymatococcus amazonicus* (Berg, 2001). *Dorstenia* was suggested to be poorly adapted for LDD (Berg & Hijman, 1999). In addition, small *Dorstenia* seeds of forest undergrowth species often germinate shortly after maturity, further reducing the chances of LDD (Berg, 2001). Low probability of LDD may explain the single origin of Neotropical *Dorstenia* and the monophyly of the two Central/North America clades. Furthermore, our results suggest a comparatively faster succession of speciation events following establishment of *Dorstenia* in the Neotropics, suggesting rapid speciation after LDD has been an important process in shaping the origin of Neotropical diversity.

Ree and Sanmartín (2018) recently raised methodological concerns with models including both anagenetic and cladogenetic processes, especially the DEC and DEC+J models. In the main analyses presented here, we focused on the results of DEC-based models (i.e., DEC m0, DEC m1, DEC+J m0, DEC+J m1). We also did reconstructions with DIVA-like and BayArea-like based models. The difference among DEC, DIVA-like and BayArea-like models lie mainly in the cladogenetic process they assume: DEC and DIVA models explore both sympatric speciation and vicariance processes while BayArea only explores sympatric speciation (Matzke, 2013b). We compared the models with the Akaike information criterion (AIC). Whether or not founder-event speciation was allowed, led the BayArea model to rank from the best to the worst model (Table S2) in both time-stratified and time-constant analyses. AIC has been argued not to be a good criterion for models including both cladogenetic and anagenetic events due to a bias of the likelihood to favor time-independent cladogenetic processes, a problem exacerbated with the introduction of founder event-speciation in the model (Ree & Sanmartín, 2018). Zero-estimate for dispersal and strong counter-intuitive unparsimonious reconstruction may be signals for this bias. These two phenomena were not observed in our reconstruction (Table S2, Figure S3). We did not rely on model selection in this study. Instead, we emphasize that all of the models we used (incl. DEC, DIVA-like and BayArea-like based models) led to very similar reconstructions (Figure 3, Figure S3).

### Taxonomic implications

Some of the currently recognized genera of Dorstenieae and all sections of *Dorstenia* may need to be modified based on the results from our analyses. Using 102 genes (132,753 bp), we obtained the same relationships among *Malaisia*, *Broussonetia* and *Allaeanthus* as a previous study based on one chloroplast and one nuclear gene (Chung et al., 2017), providing additional support for the recognition of *Allaeanthus* as a separate genus.

Both *Scyphosyce* and *Utsetela* were found to be nested in *Dorstenia*. *Scyphosyce* is a genus of two species from western Africa (Berg, 1977). We sampled both species of *Scyphosyce* in our analyses and found them sister to each other, suggesting their placement within *Dorstenia* was unlikely to be caused by misidentification. The basal grade of the *Dorstenia* s.l. clade was formed by species of sections *Nothodorstenia* and *Xylodorstenia* of the genus *Dorstenia* and by the genera *Scyphosyce* and *Utsetela,* all of which share woody habit and larger seeds, which are referred to by Berg and Hijman (1999) as macrosperms. This basal grade shares other characteristics as well. The inflorescences of most Dorstenieae genera are bisexual (some species of *Broussonetia, Allaeanthus*, and *Fatoua* have unisexual inflorescences). The macrospermous species commonly have only one to a few pistillate flowers per inflorescence which produce one to few large seeds per infructescence). The remaining species of *Dorstenia* are herbaceous and have several to numerous pistillate flowers per inflorescence which can produce numerous smaller seeds (Berg & Hijman, 1999). *Utsetela* is a genus of two species from western Africa (Berg, 1977; Jongkind, 1995). Only one species of *Utsetela* was sampled in this study. Considering the comparatively long branches of *U. gabonensis*, *D. alta*, and *D. mannii*, this relationship could be caused by a long-branch attraction artefact. To test this, we excluded the three species of *Dorstenia* which clustered with *U. gabonensis* (*D. alta*, and *D. mannii* and *D. turbinata*) and reran the phylogenetic analyses with both methods. *Utsetela gabonensis* was still in the same position (nested in *Dorstenia*) after excluding these three species (results not shown). Sampling the other species, *U. neglecta* (Jongkind, 1995), would be necessary to confirm this relationship and draw any taxonomic conclusions. Some differences among this basal grade of taxa include: tepals of individual flowers in the inflorescences of *Dorstenia* are connate, while they are free in *Scyphosyce* and *Utsetela* (Berg, 1977). While all *Scyphosyce*, *Utsetela* and *Dorstenia* have drupe(let) of fruit, their receptacles are different in shape and the filaments are far more elongated in *Utsetela* then in other two genera (Berg, 1977; Berg & Hijman, 1999). Despite some differences, similarities in woody habit, fruit type, macrospermy, and the phylogenetic reconstruction presented here suggest that merging *Scyphosyce* and *Utsetela* into *Dorstenia* may be a reasonable taxonomic outcome to preserve the monophyly of *Dorstenia*. An alternative option would be to separate those clades into two separate genera as elaborated on below.

Nine sections were proposed by Berg and Hijman (1999) in *Dorstenia* based on inflorescence, habit and life form (i.e. geophytes, phanerophytes, hemicryptophytes) characters. Our sampled species represented all the sections except *Bazzemia*, which consists of a single species in Mozambique. None of the eight sampled sections were found to be monophyletic in this study (Figure S1) and the phylogeny can help inform future subgeneric classification of *Dorstenia*. Of particular interest to consider taxonomically are the two most basal *Dorstenia* clades that include members of the sections *Xylodorstenia* and *Nothodorstenia* as well as species from two other genera (*Scyphosyce* and *Utsetela*). Misiewicz and Zerega (2012) did not include any species of *Scyphosyce* and *Utsetela,* and they found sections *Xylodorstenia* monophyletic. This is not the case in our reconstruction (Figure S1), and is likely due to our increased taxonomic sampling and use of many more genes, and our reconstructions from both ML and coalescent approaches strongly supported the non-monophyletic status of these sections. An alternative taxonomic solution to sinking *Scyphosyce* and *Utsetela* into *Dorstenia* is to include some *Dorstenia* species (most of section *Nothodorstenia* and at least one member of section *Xylodorstenia – D. angusticornis*) into the genus *Scyphosyce. Dorstenia africana*, *D. kameruniana*, *D. oligogyna*, *D. djettii* and *D. dorstenioides* (the former four all included in section *Nothodorstenia*) were once classified as genus *Craterogyne* (Lanjouw, 1935)*. Dorstenia dorstenioides*, which has been proposed as the link between sections *Xylodorstenia* and *Nothodorstenia* (Berg & Hijman, 1999), was excluded in the present study due to the low number of genes represented in our main analyses (Table S1). As *Scyphyosyce* (Baillon, 1875)is an older name than *Craterogyne, Scyphosyce* would take priority for the name of a new genus.

Regarding the clade containing *Utsetela,* some of the *Dorstenia* species in that clade were recently transferred to a new genus (*Maria*) established by Machado Vianna f. et al. (2013), comprising four species in *Dorstenia* section *Xylodorstenia* (*D. alta*, *D. angusticornis*, *D. scaphigera*, *D. turbinata*). *Maria* was later found to be a homonym and renamed *Hijmania* (Vianna Filho et al., 2016). It may be necessary to include those species into the same genus under the name *Utsetela* (Pellegrin, 1928), which has priority, but until more complete taxon sampling is completed, we do not presently propose any taxonomic changes. One of the woody African macrospermous species that warrants further attention is *D. elliptica.* It was included in section *Nothodorstenia* by Berg (1978) because it had bracts resembling other members of that section. However both Misiewicz & Zerega (2012) and the present study, found *D. elliptica* to be sister to all Neotropical *Dorstenia.*

Among herbaceous species of *Dorstenia*, the presence of bracts on receptacles has traditionally been used to distinguish among sections *Emygdioa*, *Dorstenia*, *Lecanium* on the one hand and sections *Acauloma*, *Bazzemia*, *Lomatophora*, *Kosaria* on the other hand (Berg & Hijman, 1999). Although none of these sections were found to be monophyletic, the Neotropical clade contains all of the species with bracteate receptacles except *D. picta* (Figure S1). Our results suggest that traditional morphological characters for sectional delimitation within *Dorstenia* do not hold up to molecular phylogenetic scrutiny, that a close examination of alternative characters is needed, and a new intrageneric classification is warranted.

## CONCLUSION

The targeted enrichment sequencing strategy, paired with the HybPiper pipeline, proved to be an effective approach at reconstructing phylogenetic relationships in Dorstenieae using herbarium specimens. Further molecular and morphological work will be required before solving some of the taxonomic issues highlighted in this study, such as sinking *Utsetela* and *Scyphosyce* into *Dorstenia* or separating some *Dorstenia* species into either the genera of *Utsetela* or *Scyphosyce*. Dorstenieae as a whole may have originated in the joint area of continental Africa, Madagascar and Asia-Oceania area, followed by at least two independent colonizations of South America (*Brosimum* s.l. and *Dorstenia* s.l.). Some species in these two clades further dispersed to Central/North America. The mechanical processes for the long-distance dispersal of species of Dorstenieae remains an enigma. More studies on pollination and dispersal in this tribe will be required to further elucidate the biogeographic history of this group. The development of new biogeographic models and new model selection procedures will also be essential to help to clear the biogeographic history of Dorstenieae and other pantropically distributed lineages. The robust and most densely sampled Dorstenieae phylogeny presented here will be a valuable resource for further studies on character evolution in this fascinating tribe and will assist with future taxonomic revisionary work.

## ACKNOWLEDGEMENTS

We thank the Field Museum (F), the Missouri Botanical Garden (MO), the New York Botanical Garden (NY), and the Muséum national d’Histoire naturelle (P) for sharing herbarium material; Tracy Misiewicz and Caroline Storer for assistance with DNA extraction; Charles Foster for providing helpful advice on running MCMCTree; and Santiago Ramirez-Barahona for information on fossil calibrations. This work was supported by a China Scholarship Council (CSC) PhD grant (grant No. 201506140077) and a research grant from the International Association for Plant Taxonomy (IAPT) to Q.Z. Sequencing costs were covered by Agence Nationale de la Recherche grant ANR-12-JVS7-0015-01 to H.S.

## SUPPLEMENTARY INFORMATION

**Figure S1.** Maximum Likelihood tree (a) and species tree (b) with bootstrap support value and tip names (including section names in *Dorstenia*)

**Figure S2.** Divergence time estimate by MCMCTree with two different prior settings and r8s: combined results of two independent runs with prior set1 (a) and set2 (b) by MCMCTree and r8s (c)

**Figure S3.** Biogeographic reconstruction with unconstrained (model 0) or constrained (model 1) models with DEC, DIVA-like and BayArea-like based models with internal nodes labelled with discrete states: a-c, DEC; d, illustrated with pie chart on internal nodes for the result of time-stratified DEC+J model; e-h, Bayarea-like; i-l, DIVA-like model with model 0 and 1, with or without founder-event speciation process (+J), detail of DEC+J model with nodes labelled with discrete states see Figure 3.

**Table S1.** List of specimens collected in this study.

**Table S2.**
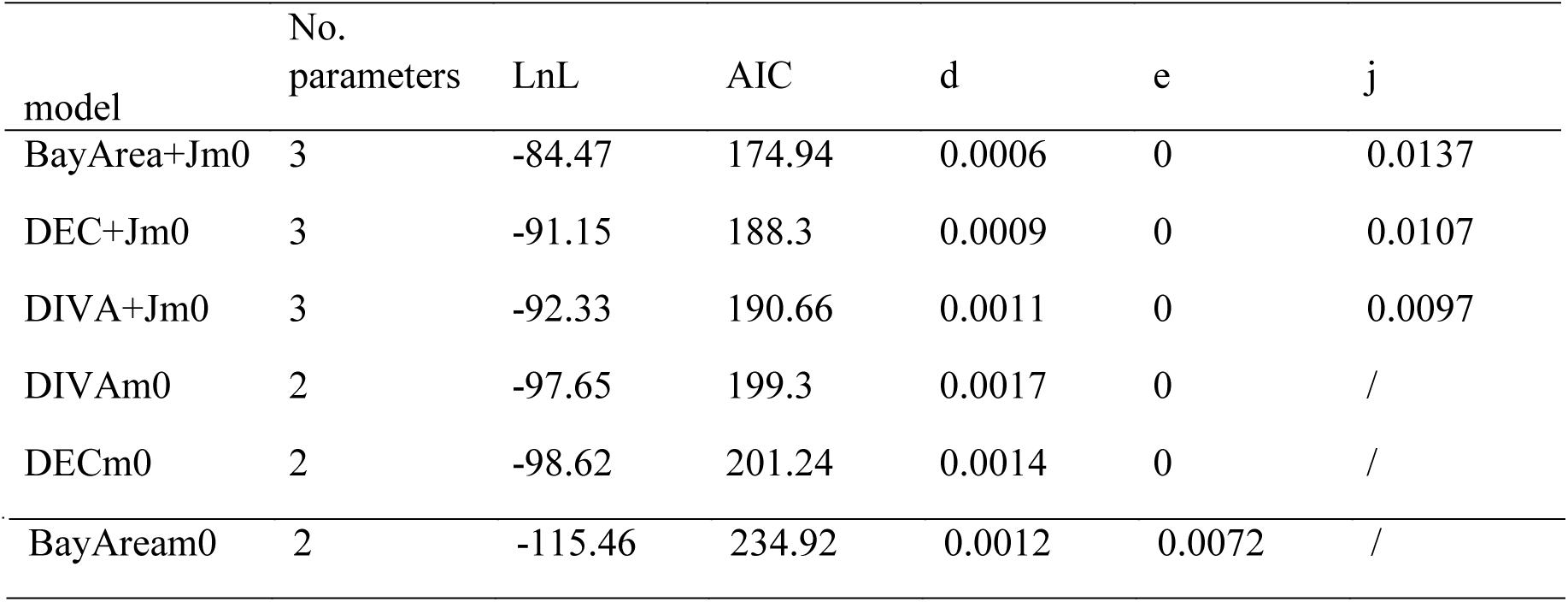
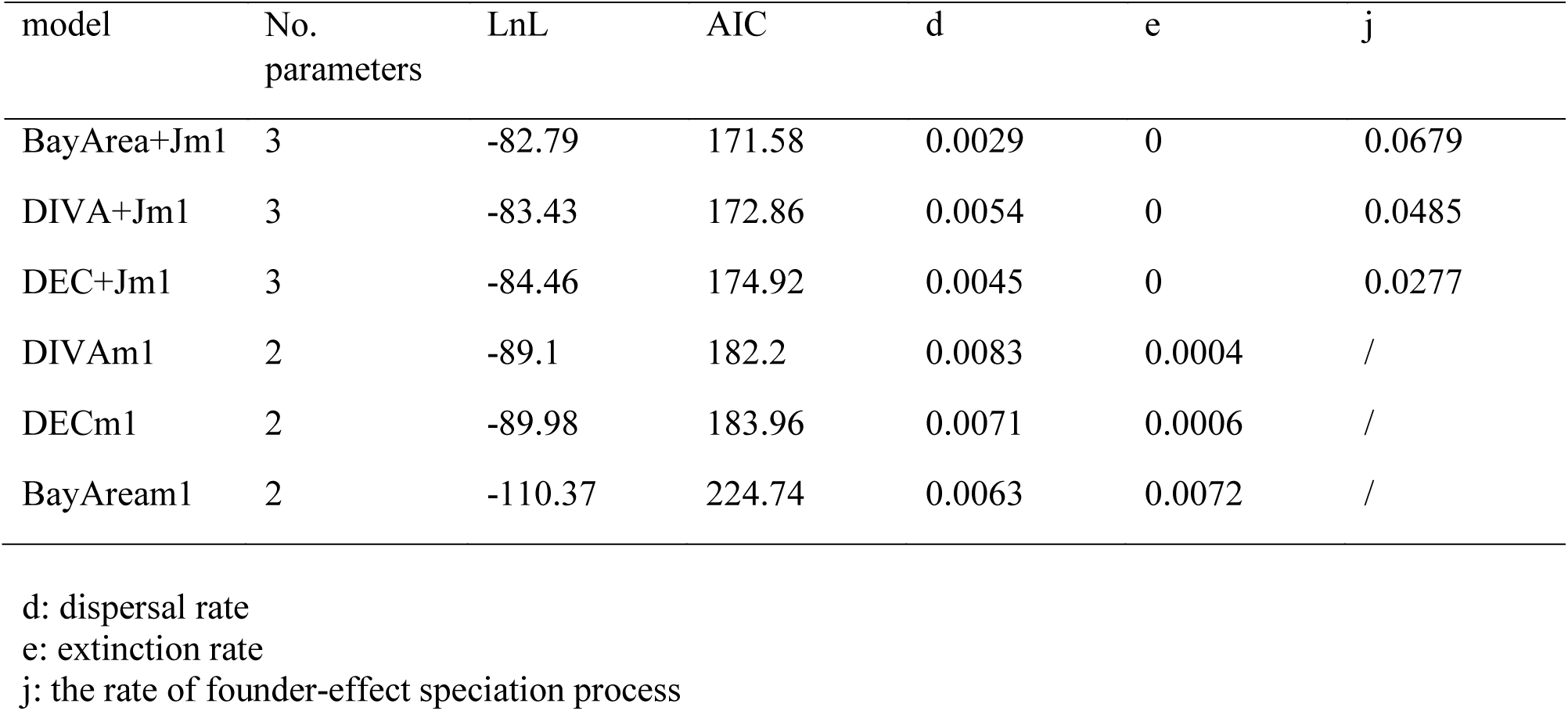
AIC of biogeographic reconstruction with time-constant (model0, a) or time-stratified (model1, b) DEC, DIVA-like and BayArea-like based models.

**Table S3.**
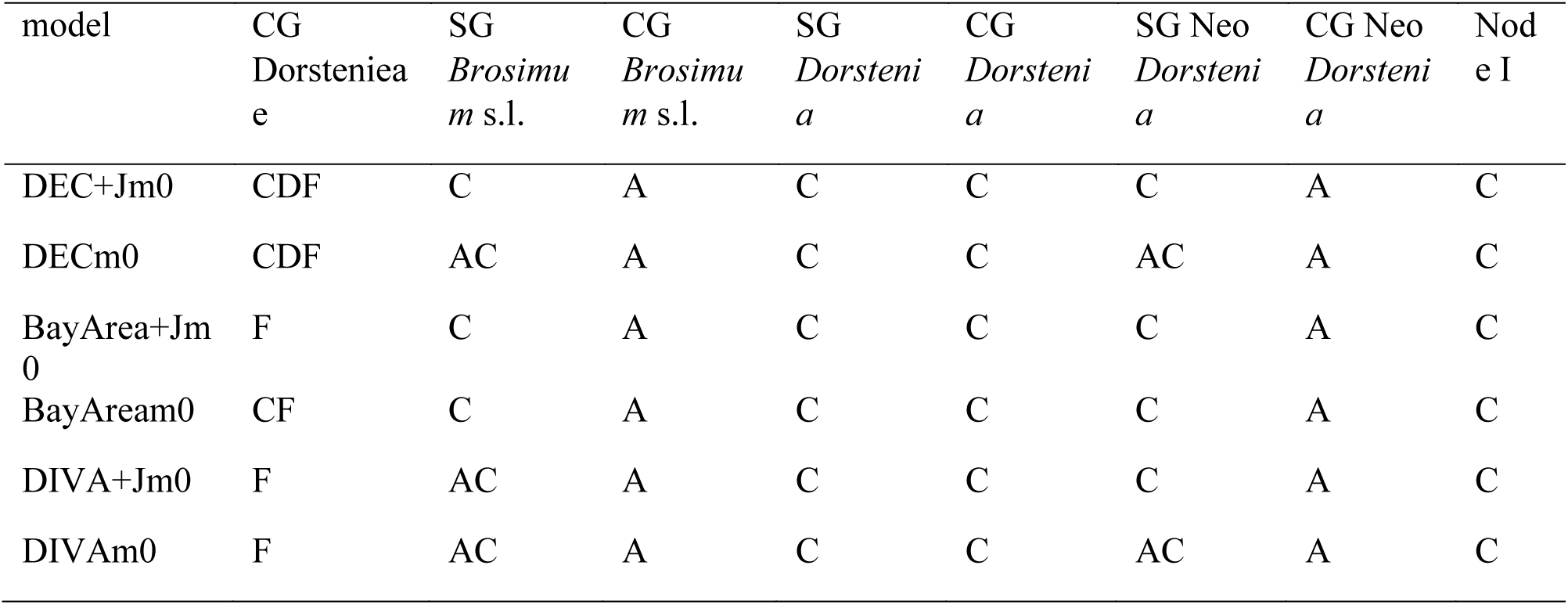
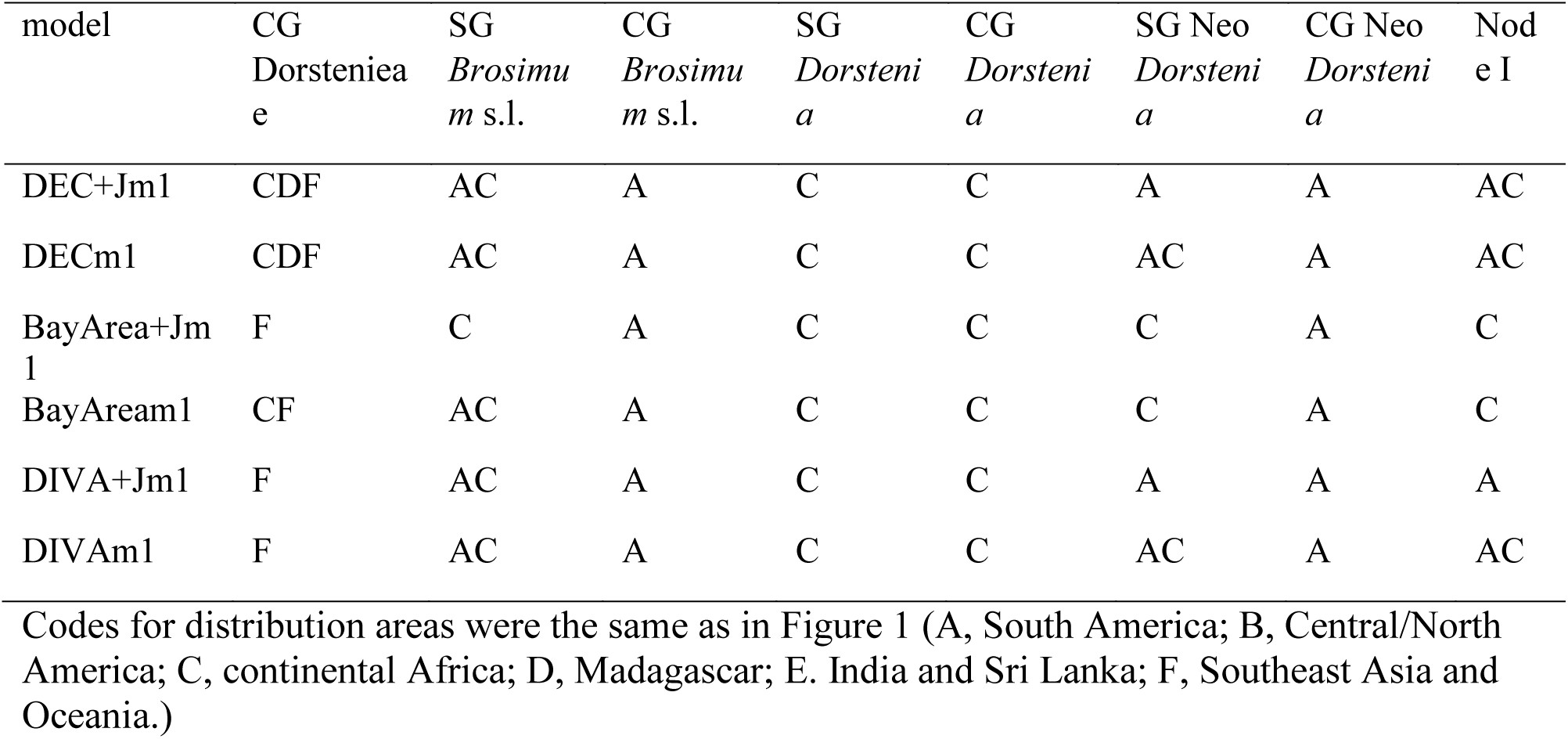
List of ancestral distribution area estimated for several nodes by BioGeoBEARS. a) time-constant models (model0); b) time-stratified models (model1)

