## Supplementary figures and images for "Long-distance dispersal shaped the diversity of tribe Dorstenieae (Moraceae)"

### Figure S1

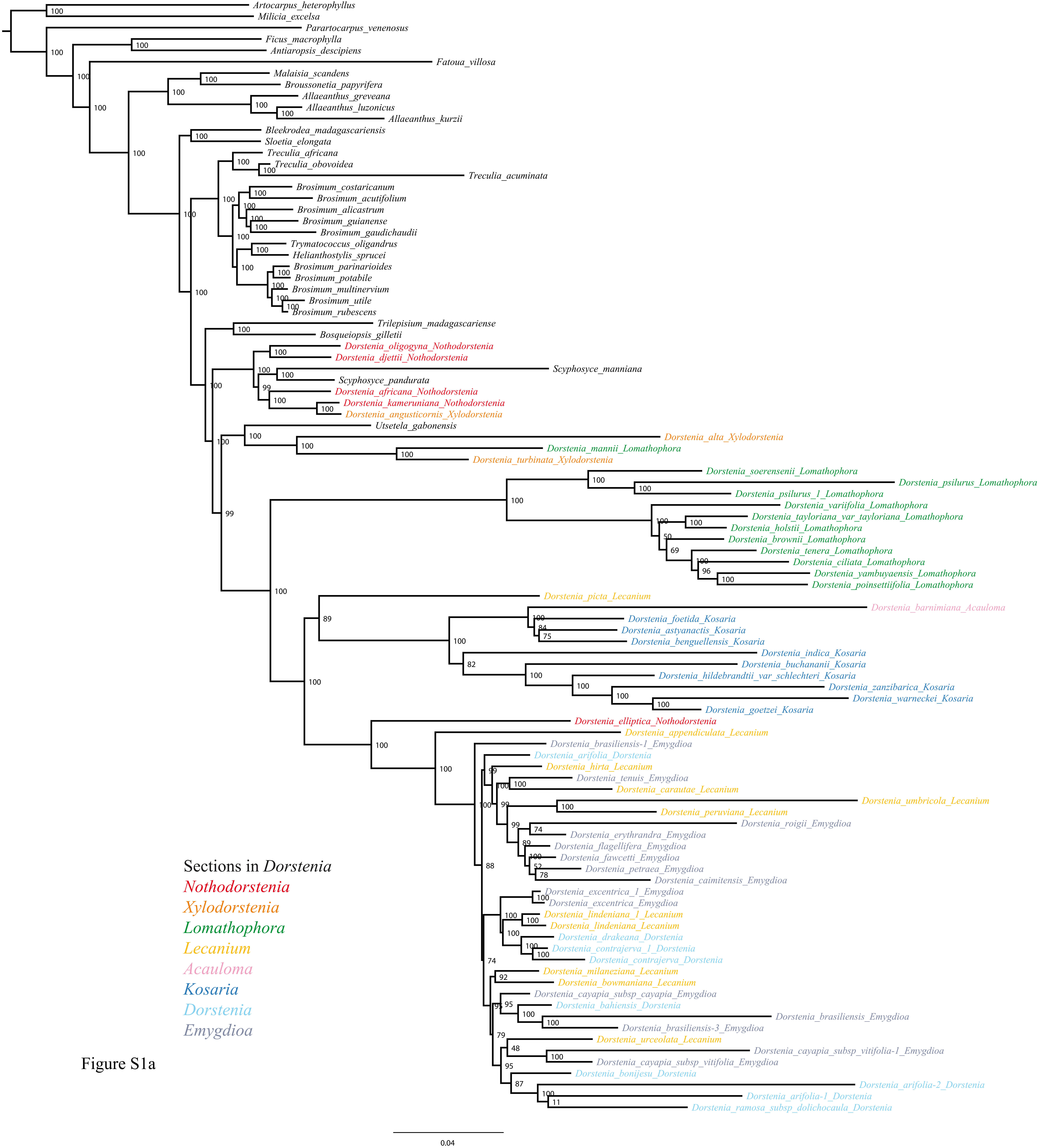

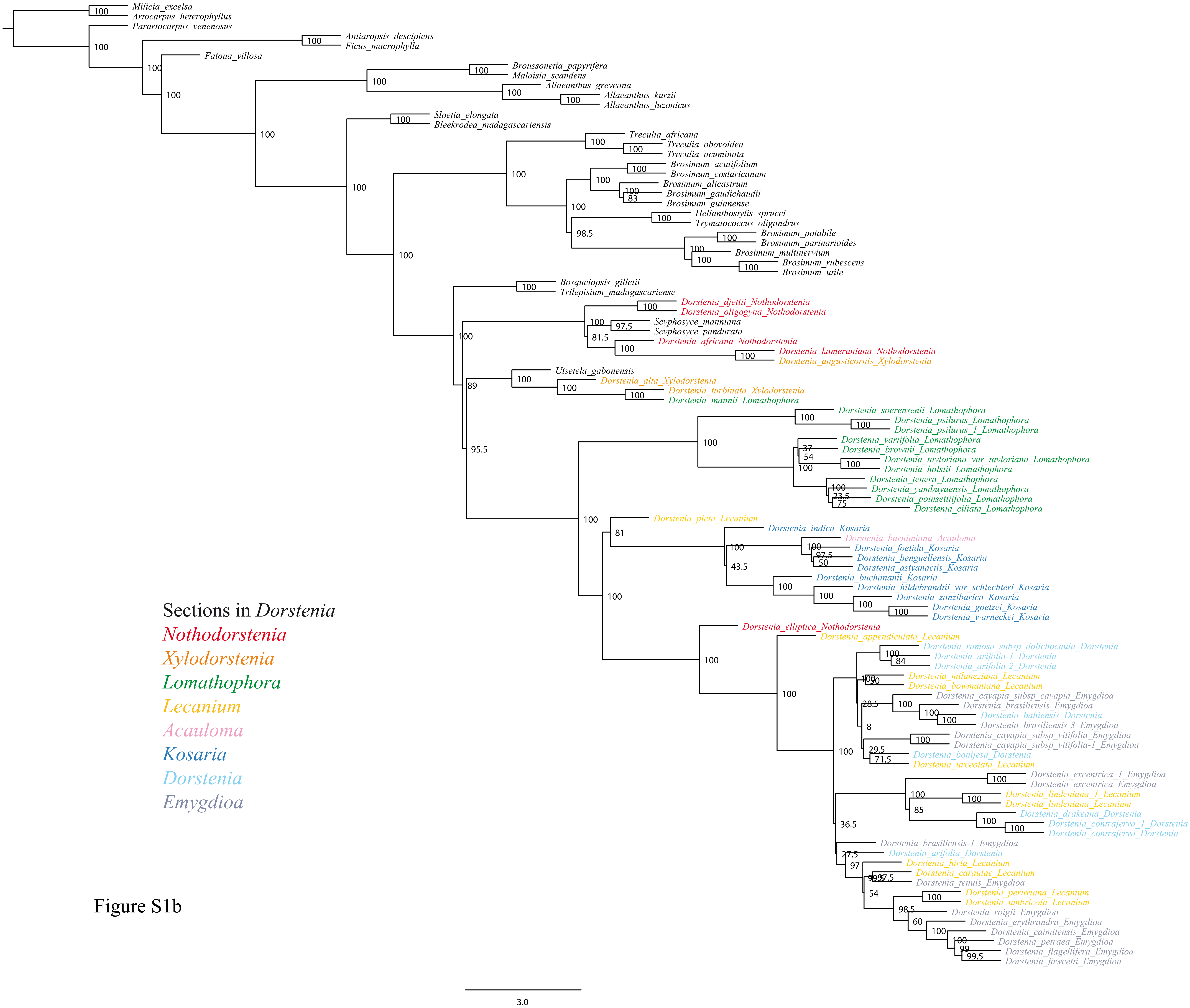

### Figure S2

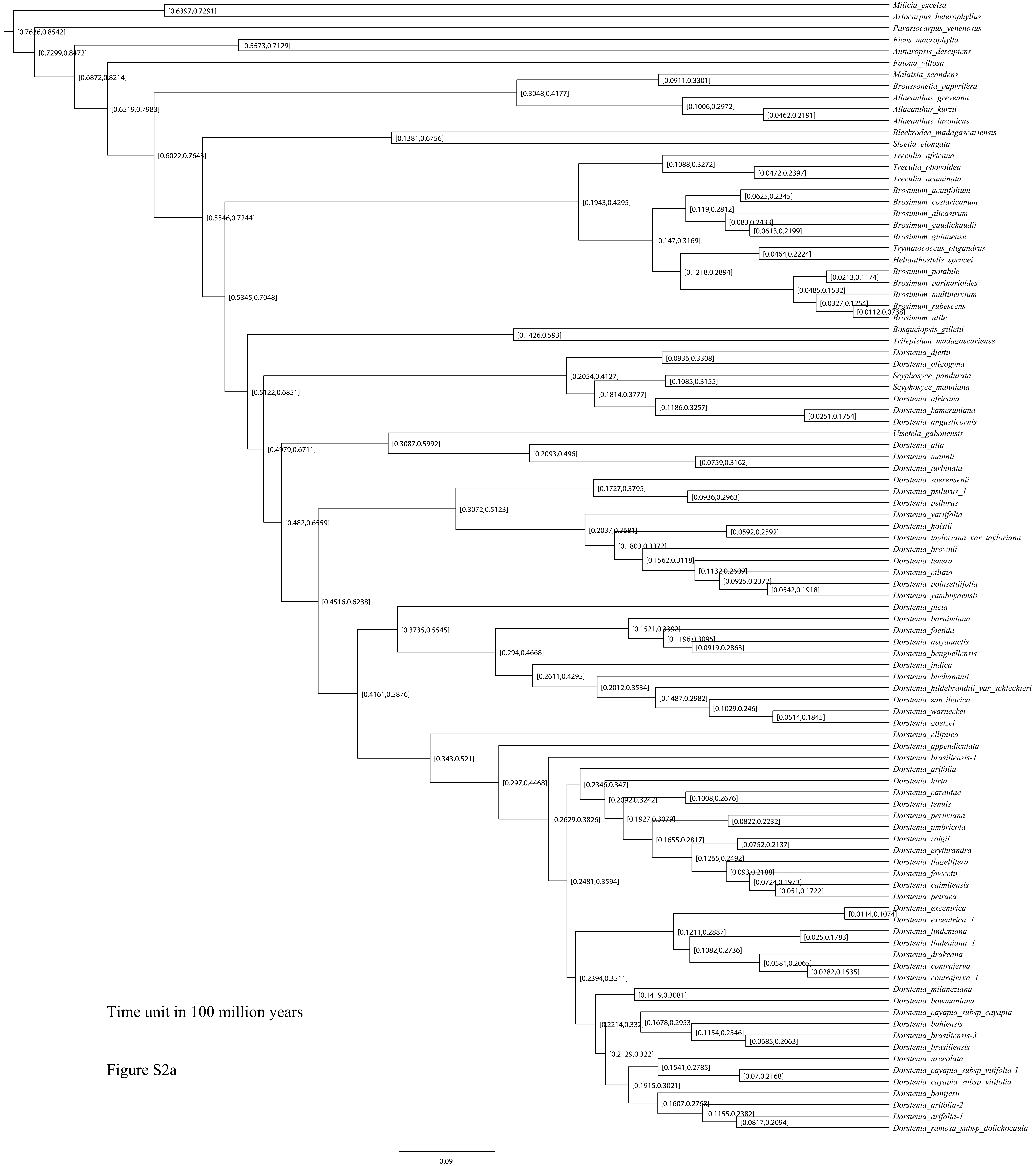

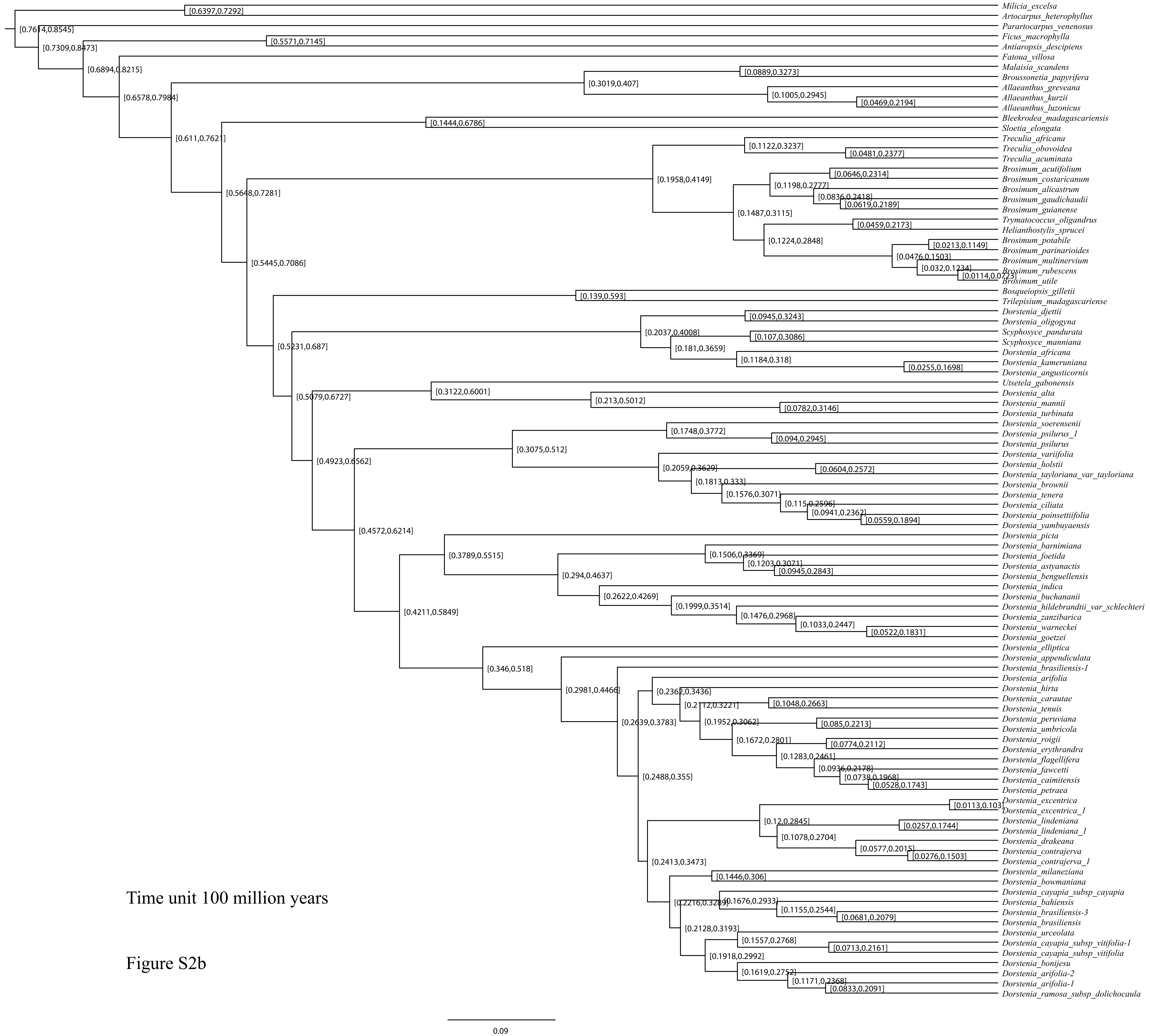

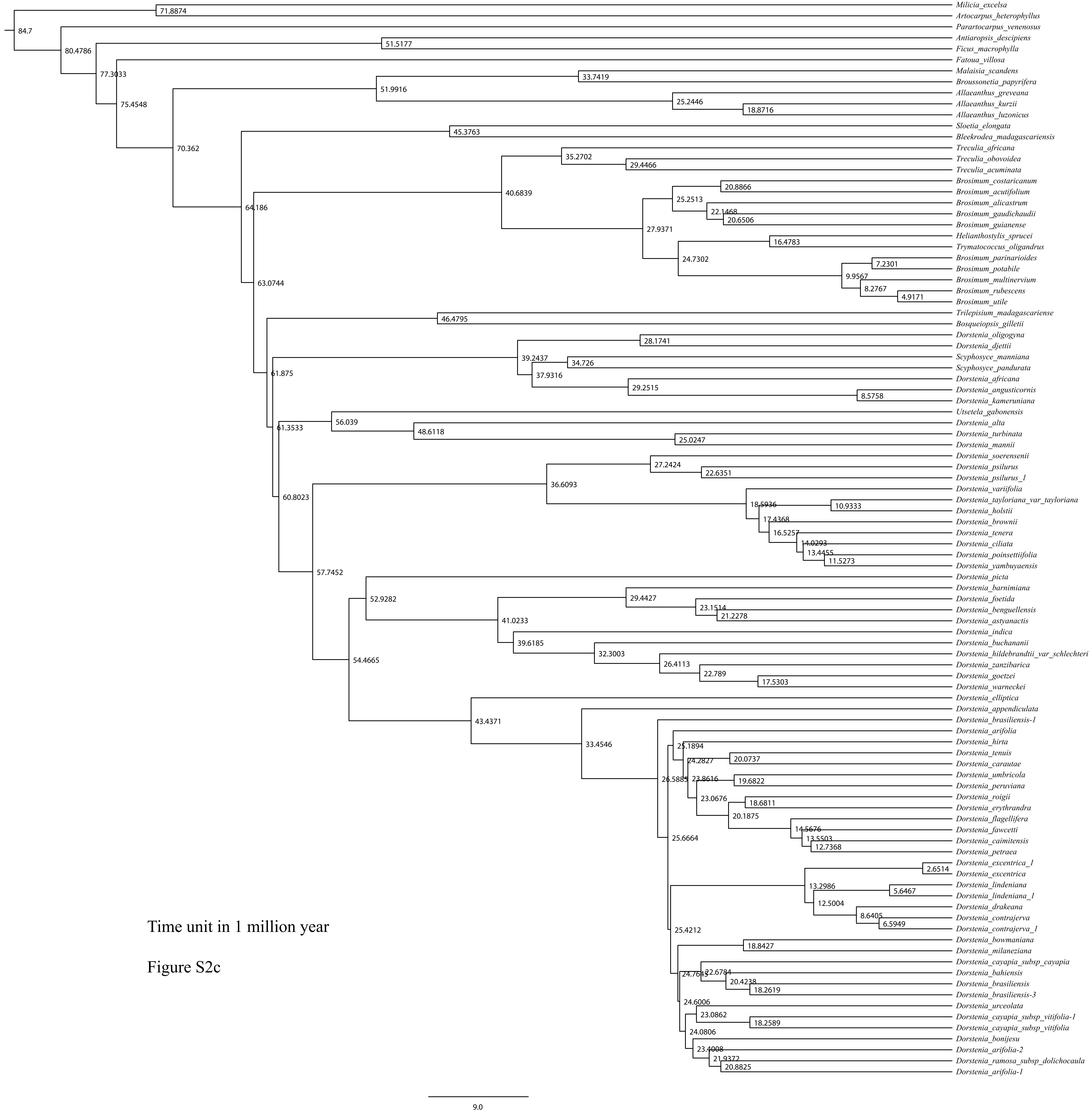
