## Supplementary material for "Long-distance dispersal shaped the diversity of tribe Dorstenieae (Moraceae)": Figure S3

**DEC model0 on Dorstenieae**
**ancstates: global optim, 3 areas max. d=0.0014; e=0; j=0; LnL=−98.62**

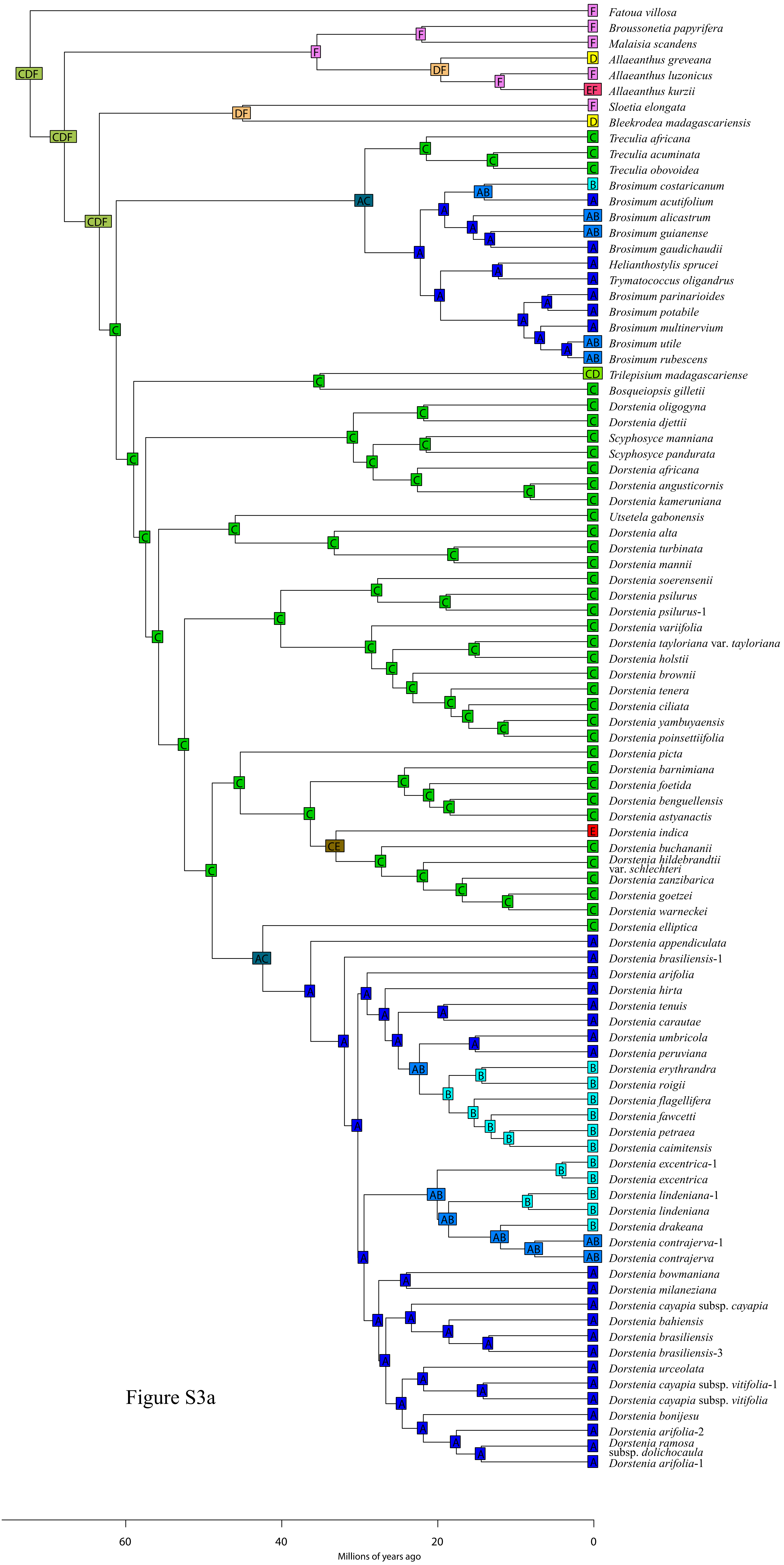


Figure S3a

**DEC+J model0 on Dorstenieae**
**ancstates: global optim, 3 areas max. d=9e−04; e=0; j=0.0107; LnL=−91.15**

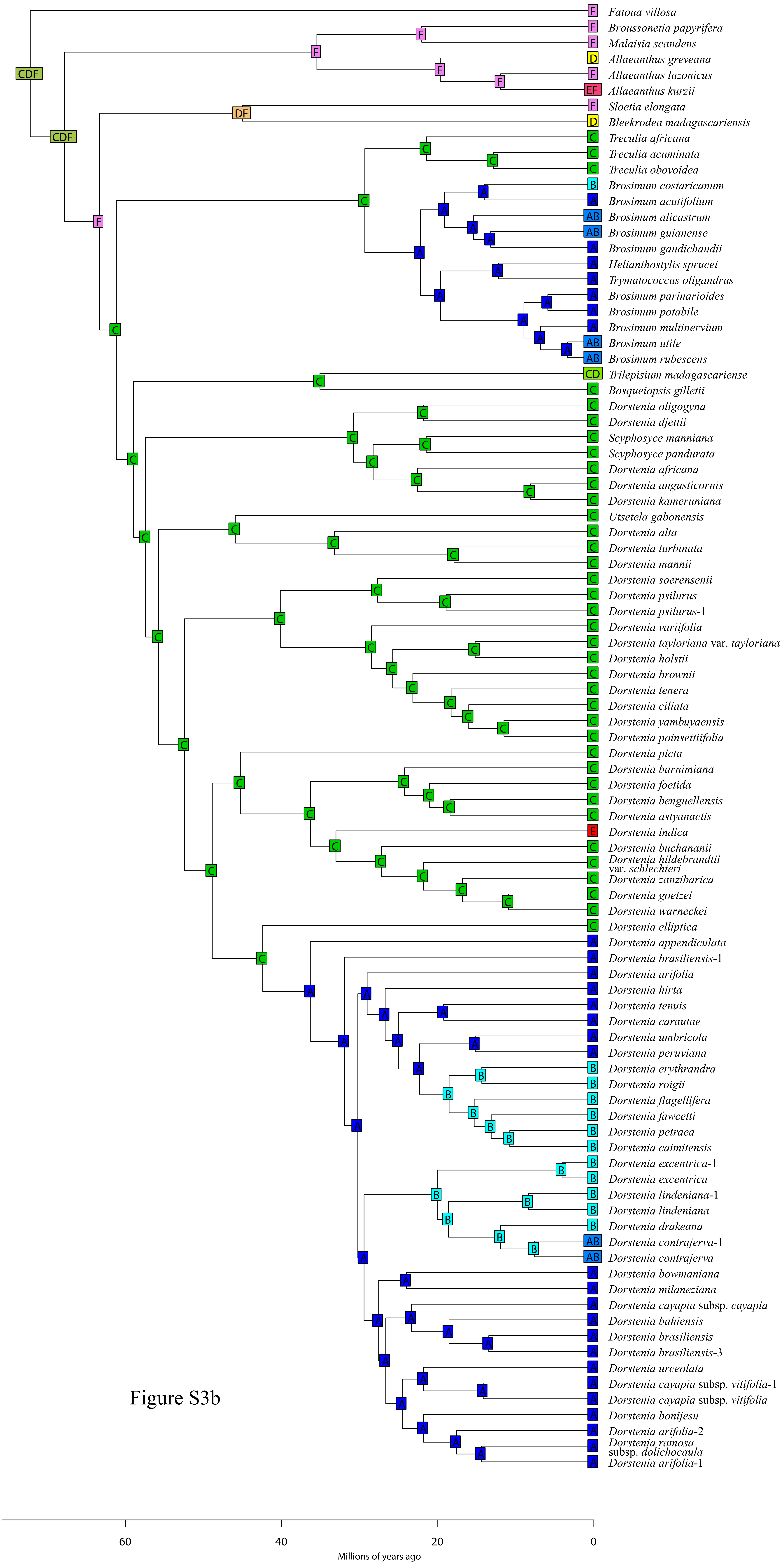

**DEC model1 on Dorstenieae**

**ancstates: global optim, 3 areas max. d=0.0071; e=6e−04; j=0; LnL=−89.98**

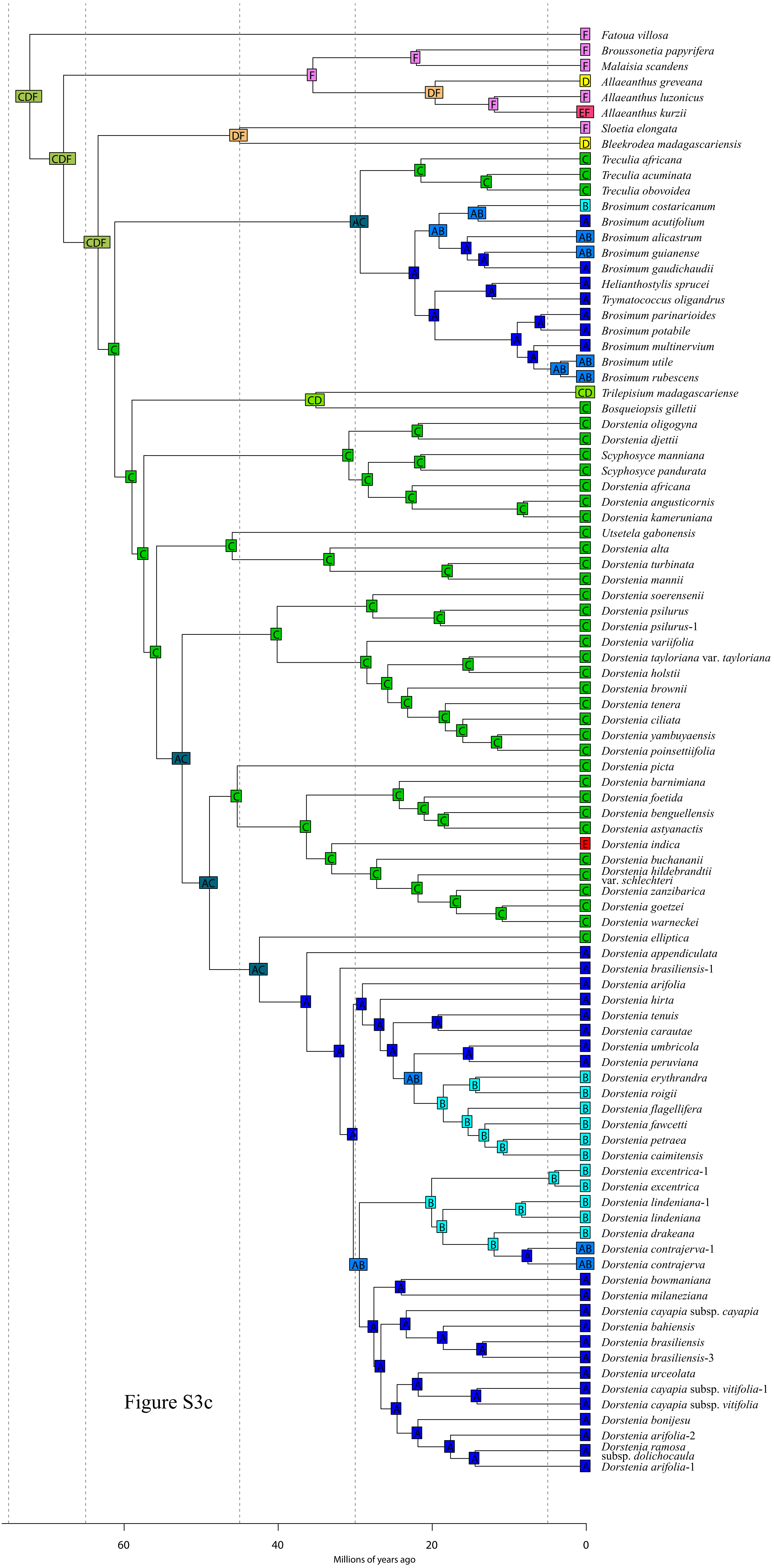


Figure S3c

**DEC+J model1 on Dorstenieae**
**ancstates: global optim, 3 areas max. d=0.0045; e=0; j=0.0277; LnL=−84.46**

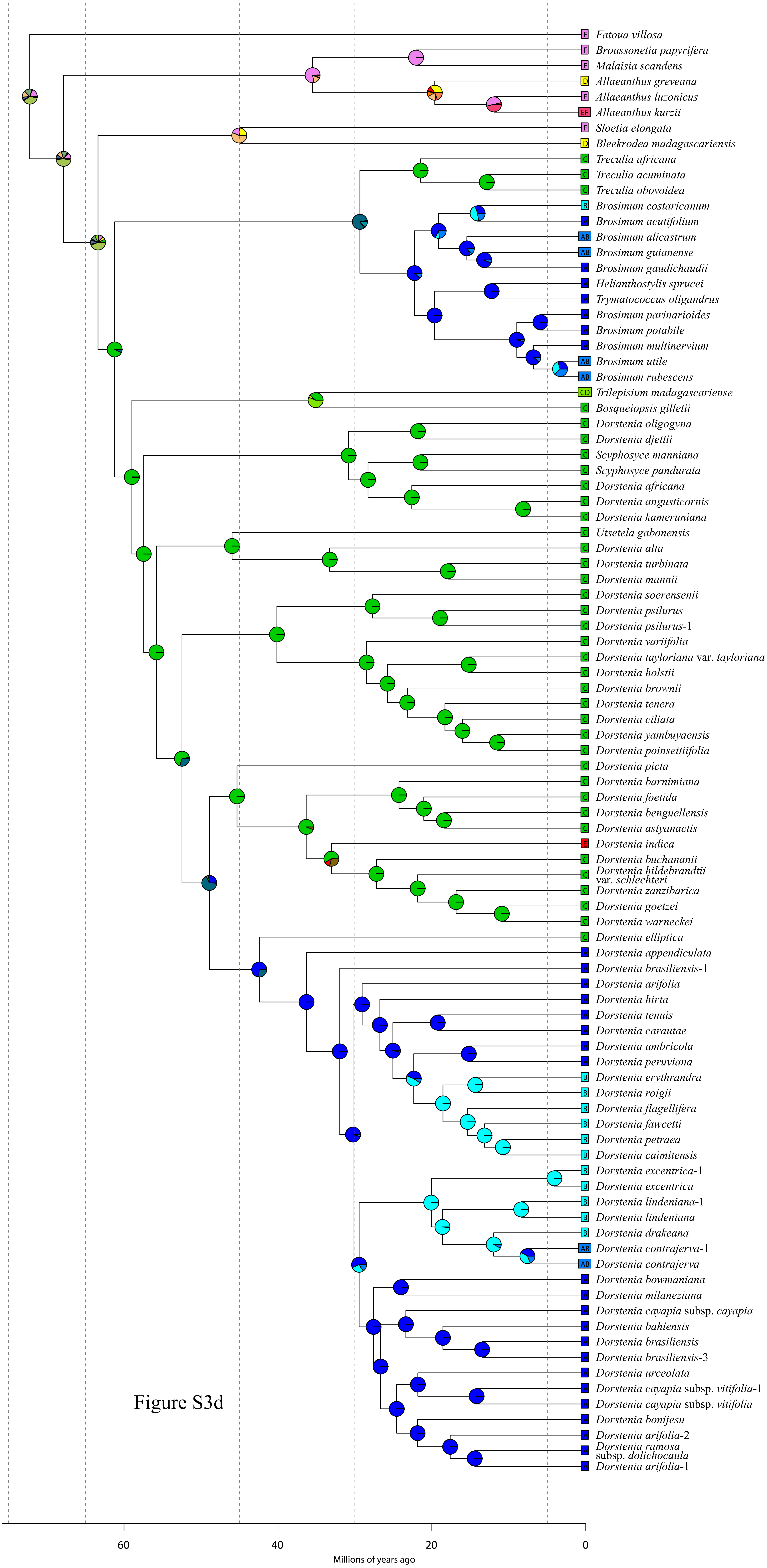


Figure S3d

**BayArea model0 on Dorstenieae**
**ancstates: global optim, 3 areas max. d=0.0012; e=0.0072; j=0; LnL=−115.46**

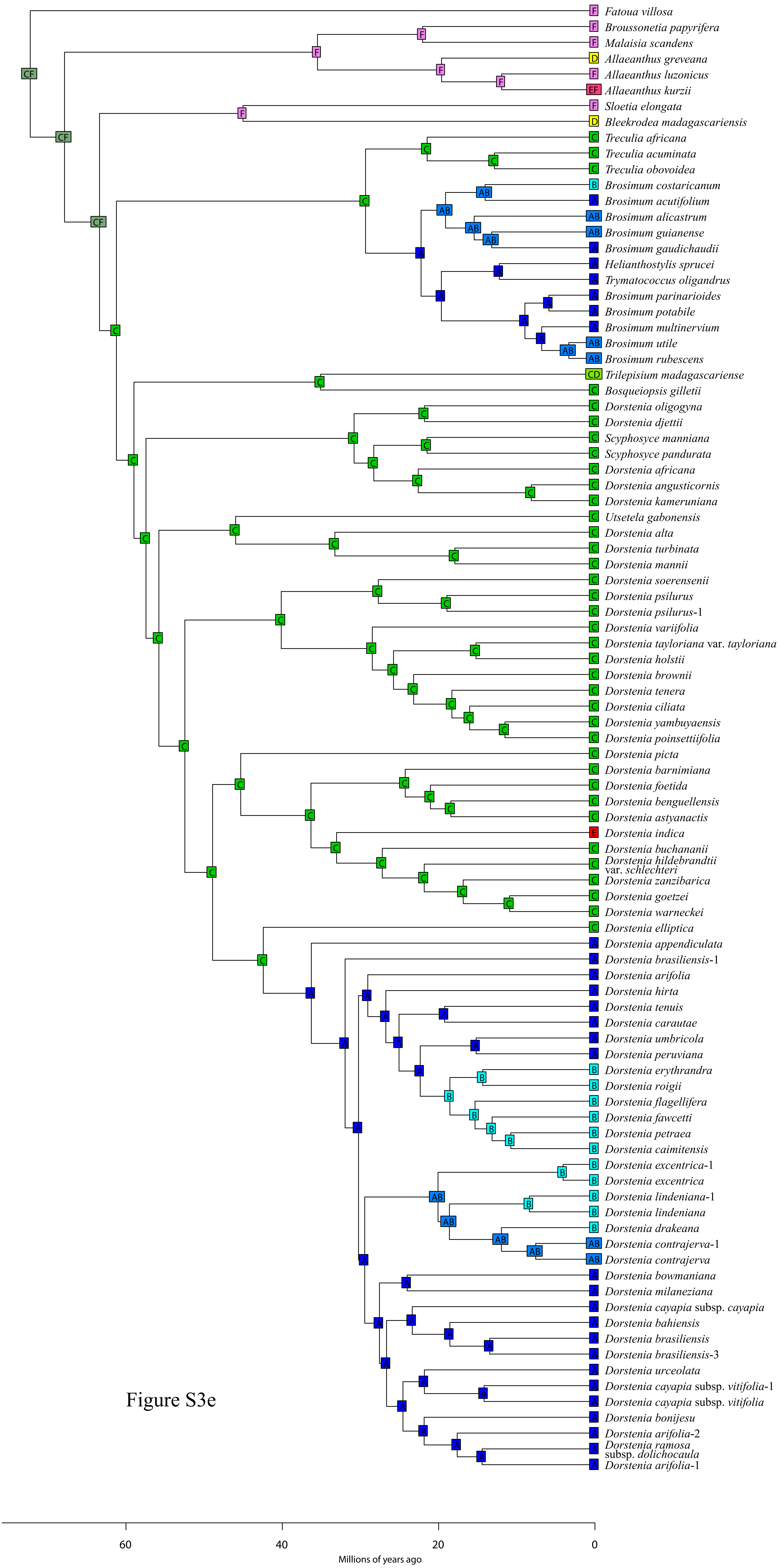


Figure S3e

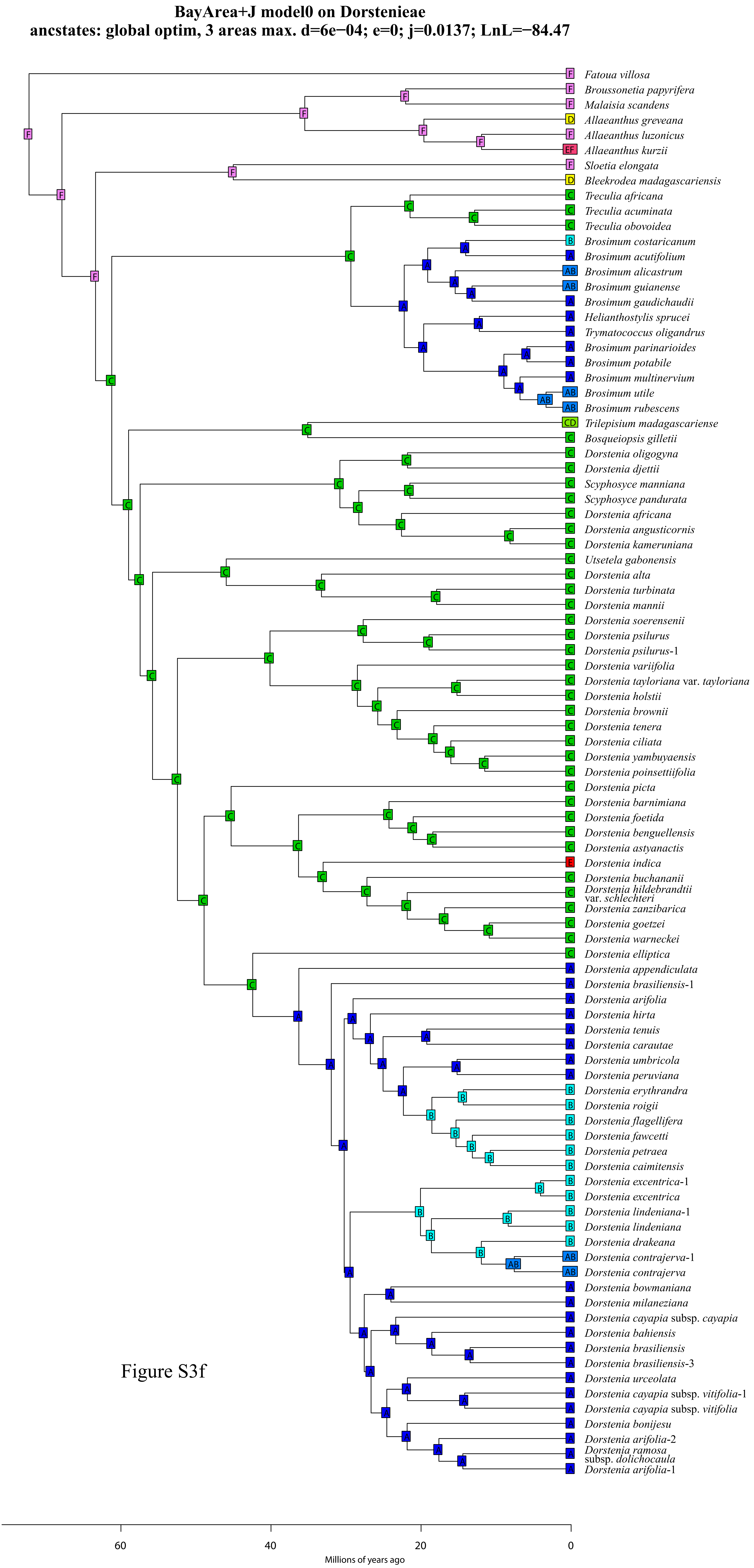
BayArea+J model0 on Dorstenieae
ancstates: global optim, 3 areas max. d=6e−04; e=0; j=0.0137; LnL=−84.47
Fatoua villosa
Broussonetia papyrifera
Malaisia scandens
Allaeanthus greveana
Allaeanthus luzonicus
Allaeanthus kurzii
Sloetia elongata
Bleekrodea madagascariensis
Treculia africana
Treculia acuminata
Treculia obovoidea
Brosimum costaricanum
Brosimum acutifolium
Brosimum alicastrum
Brosimum guianense
Brosimum gaudichaudii
Helianthostylis sprucei
Trymatococcus oligandrus
Brosimum parinarioides
Brosimum potabile
Brosimum multinervium
Brosimum utile
Brosimum rubescens
Trilepisium madagascariense
Bosqueiopsis gilletii
Dorstenia oligogyna
Dorstenia djettii
Scyphosyce manniana
Scyphosyce pandurata
Dorstenia africana
Dorstenia angusticornis
Dorstenia kameruniana
Utsetela gabonensis
Dorstenia alta
Dorstenia turbinata
Dorstenia mannii
Dorstenia soerensenii
Dorstenia psilurus
Dorstenia psilurus-1
Dorstenia variifolia
Dorstenia tayloriana var. tayloriana
Dorstenia holstii
Dorstenia brownii
Dorstenia tenera
Dorstenia ciliata
Dorstenia yambuyaensis
Dorstenia poinsettiifolia
Dorstenia picta
Dorstenia barnimiana
Dorstenia foetida
Dorstenia benguellensis
Dorstenia astyanactis
Dorstenia indica
Dorstenia buchananii
Dorstenia hildebrandtii var. schlechteri
Dorstenia zanzibarica
Dorstenia goetzei
Dorstenia warneckei
Dorstenia elliptica
Dorstenia appendiculata
Dorstenia brasiliensis-1
Dorstenia arifolia
Dorstenia hirta
Dorstenia tenuis
Dorstenia carautae
Dorstenia umbricola
Dorstenia peruviana
Dorstenia erythrandra
Dorstenia roigii
Dorstenia flagellifera
Dorstenia fawcetti
Dorstenia petraea
Dorstenia caimitensis
Dorstenia excentrica-1
Dorstenia excentrica
Dorstenia lindeniana-1
Dorstenia lindeniana
Dorstenia drakeana
Dorstenia contrajerva-1
Dorstenia contrajerva
Dorstenia bowmaniana
Dorstenia milaneziana
Dorstenia cayapia subsp. cayapia
Dorstenia bahiensis
Dorstenia brasiliensis
Dorstenia brasiliensis-3
Dorstenia urceolata
Dorstenia cayapia subsp. vitifolia-1
Dorstenia cayapia subsp. vitifolia
Dorstenia bonijesu
Dorstenia arifolia-2
Dorstenia ramosa subsp. dolichocaula
Dorstenia arifolia-1
60
40
20
0
Millions of years ago

Figure S3f

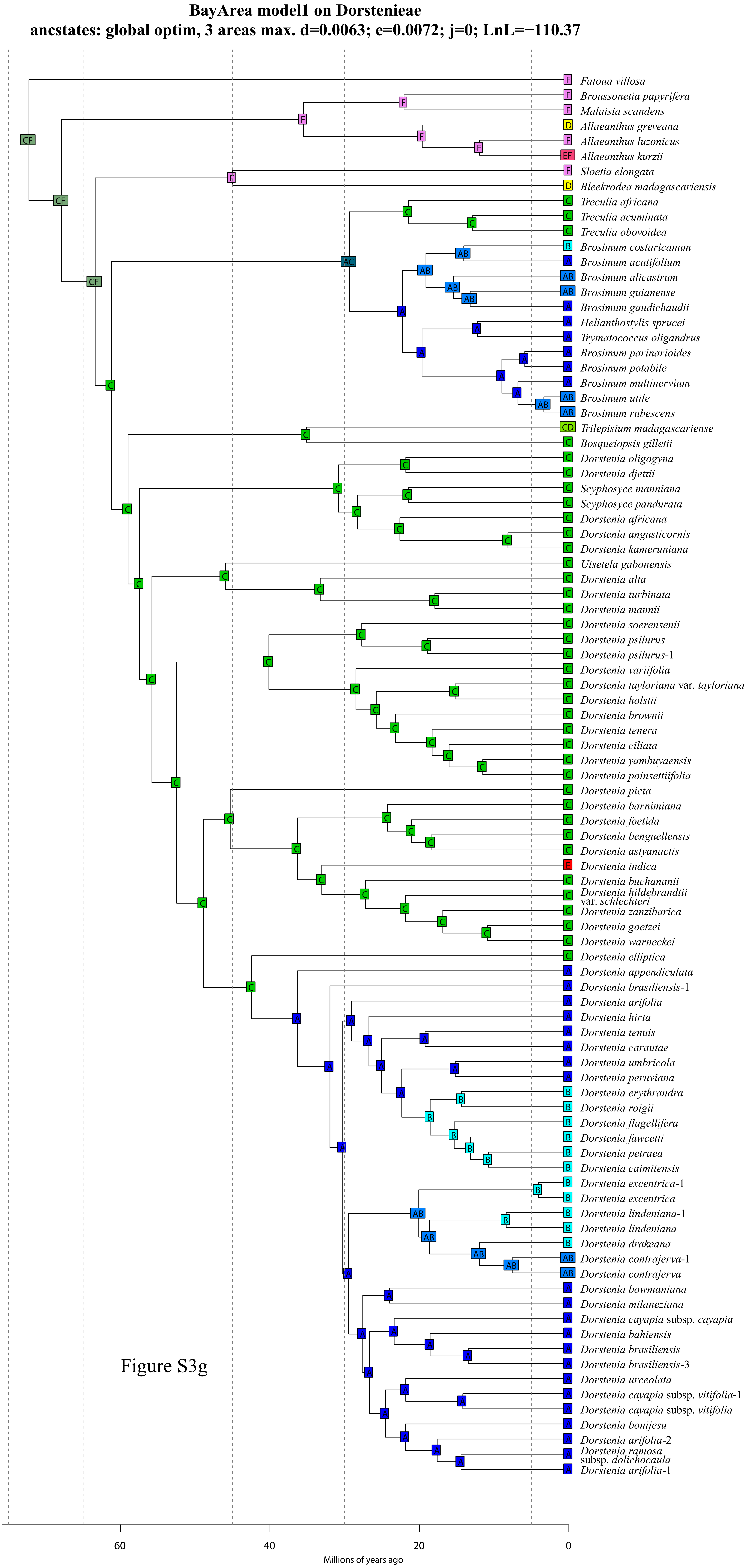
BayArea model1 on Dorstenieae
ancstates: global optim, 3 areas max. d=0.0063; e=0.0072; j=0; LnL=−110.37
Fatoua villosa
Broussonetia papyrifera
Malaisia scandens
Allaeanthus greveana
Allaeanthus luzonicus
Allaeanthus kurzii
Sloetia elongata
Bleekrodea madagascariensis
Treculia africana
Treculia acuminata
Treculia obovoidea
Brosimum costaricanum
Brosimum acutifolium
Brosimum alicastrum
Brosimum guianense
Brosimum gaudichaudii
Helianthostylis sprucei
Trymatococcus oligandrus
Brosimum parinarioides
Brosimum potabile
Brosimum multinervium
Brosimum utile
Brosimum rubescens
Trilepisium madagascariense
Bosqueiopsis gilletii
Dorstenia oligogyna
Dorstenia djettii
Scyphosyce manniana
Scyphosyce pandurata
Dorstenia africana
Dorstenia angusticornis
Dorstenia kameruniana
Utsetela gabonensis
Dorstenia alta
Dorstenia turbinata
Dorstenia mannii
Dorstenia soerensenii
Dorstenia psilurus
Dorstenia psilurus-1
Dorstenia variifolia
Dorstenia tayloriana var. tayloriana
Dorstenia holstii
Dorstenia brownii
Dorstenia tenera
Dorstenia ciliata
Dorstenia yambuyaensis
Dorstenia poinsettiifolia
Dorstenia picta
Dorstenia barnimiana
Dorstenia foetida
Dorstenia benguellensis
Dorstenia astyanactis
Dorstenia indica
Dorstenia buchananii
Dorstenia hildebrandtii var. schlechteri
Dorstenia zanzibarica
Dorstenia goetzei
Dorstenia warneckei
Dorstenia elliptica
Dorstenia appendiculata
Dorstenia brasiliensis-1
Dorstenia arifolia
Dorstenia hirta
Dorstenia tenuis
Dorstenia carautae
Dorstenia umbricola
Dorstenia peruviana
Dorstenia erythrandra
Dorstenia roigii
Dorstenia flagellifera
Dorstenia fawcetti
Dorstenia petraea
Dorstenia caimitensis
Dorstenia excentrica-1
Dorstenia excentrica
Dorstenia lindeniana-1
Dorstenia lindeniana
Dorstenia drakeana
Dorstenia contrajerva-1
Dorstenia contrajerva
Dorstenia bowmaniana
Dorstenia milaneziana
Dorstenia cayapia subsp. cayapia
Dorstenia bahiensis
Dorstenia brasiliensis
Dorstenia brasiliensis-3
Dorstenia urceolata
Dorstenia cayapia subsp. vitifolia-1
Dorstenia cayapia subsp. vitifolia
Dorstenia bonijesu
Dorstenia arifolia-2
Dorstenia ramosa subsp. dolichocaula
Dorstenia arifolia-1
60
40
20
0
Millions of years ago

Figure S3g

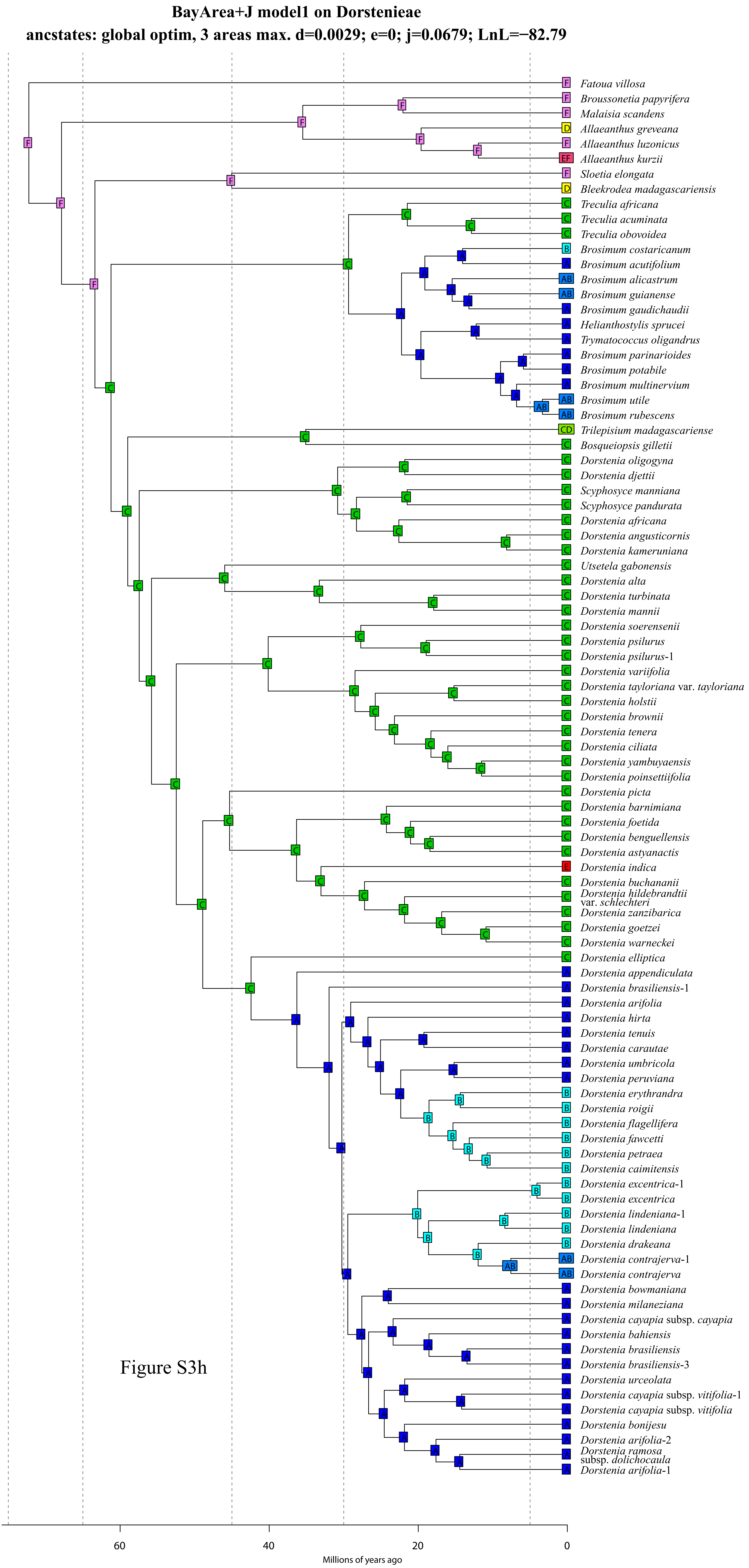
BayArea+J model1 on Dorstenieae
ancstates: global optim, 3 areas max. d=0.0029; e=0; j=0.0679; LnL=−82.79
Fatoua villosa
Broussonetia papyrifera
Malaisia scandens
Allaeanthus greveana
Allaeanthus luzonicus
Allaeanthus kurzii
Sloetia elongata
Bleekrodea madagascariensis
Treculia africana
Treculia acuminata
Treculia obovoidea
Brosimum costaricanum
Brosimum acutifolium
Brosimum alicastrum
Brosimum guianense
Brosimum gaudichaudii
Helianthostylis sprucei
Trymatococcus oligandrus
Brosimum parinarioides
Brosimum potabile
Brosimum multinervium
Brosimum utile
Brosimum rubescens
Trilepisium madagascariense
Bosqueiopsis gilletii
Dorstenia oligogyna
Dorstenia djettii
Scyphosyce manniana
Scyphosyce pandurata
Dorstenia africana
Dorstenia angusticornis
Dorstenia kameruniana
Utsetela gabonensis
Dorstenia alta
Dorstenia turbinata
Dorstenia mannii
Dorstenia soerensenii
Dorstenia psilurus
Dorstenia psilurus-1
Dorstenia variifolia
Dorstenia tayloriana var. tayloriana
Dorstenia holstii
Dorstenia brownii
Dorstenia tenera
Dorstenia ciliata
Dorstenia yambuyaensis
Dorstenia poinsettiifolia
Dorstenia picta
Dorstenia barnimiana
Dorstenia foetida
Dorstenia benguellensis
Dorstenia astyanactis
Dorstenia indica
Dorstenia buchananii
Dorstenia hildebrandtii var. schlechteri
Dorstenia zanzibarica
Dorstenia goetzei
Dorstenia warneckei
Dorstenia elliptica
Dorstenia appendiculata
Dorstenia brasiliensis-1
Dorstenia arifolia
Dorstenia hirta
Dorstenia tenuis
Dorstenia carautae
Dorstenia umbricola
Dorstenia peruviana
Dorstenia erythrandra
Dorstenia roigii
Dorstenia flagellifera
Dorstenia fawcetti
Dorstenia petraea
Dorstenia caimitensis
Dorstenia excentrica-1
Dorstenia excentrica
Dorstenia lindeniana-1
Dorstenia lindeniana
Dorstenia drakeana
Dorstenia contrajerva-1
Dorstenia contrajerva
Dorstenia bowmaniana
Dorstenia milaneziana
Dorstenia cayapia subsp. cayapia
Dorstenia bahiensis
Dorstenia brasiliensis
Dorstenia brasiliensis-3
Dorstenia urceolata
Dorstenia cayapia subsp. vitifolia-1
Dorstenia cayapia subsp. vitifolia
Dorstenia bonijesu
Dorstenia arifolia-2
Dorstenia ramosa subsp. dolichocaula
Dorstenia arifolia-1
60
40
20
0
Millions of years ago

Figure S3h

**DIVA model0 on Dorstenieae**

**ancstates: global optim, 3 areas max. d=0.0017; e=0; j=0; LnL=−97.65**

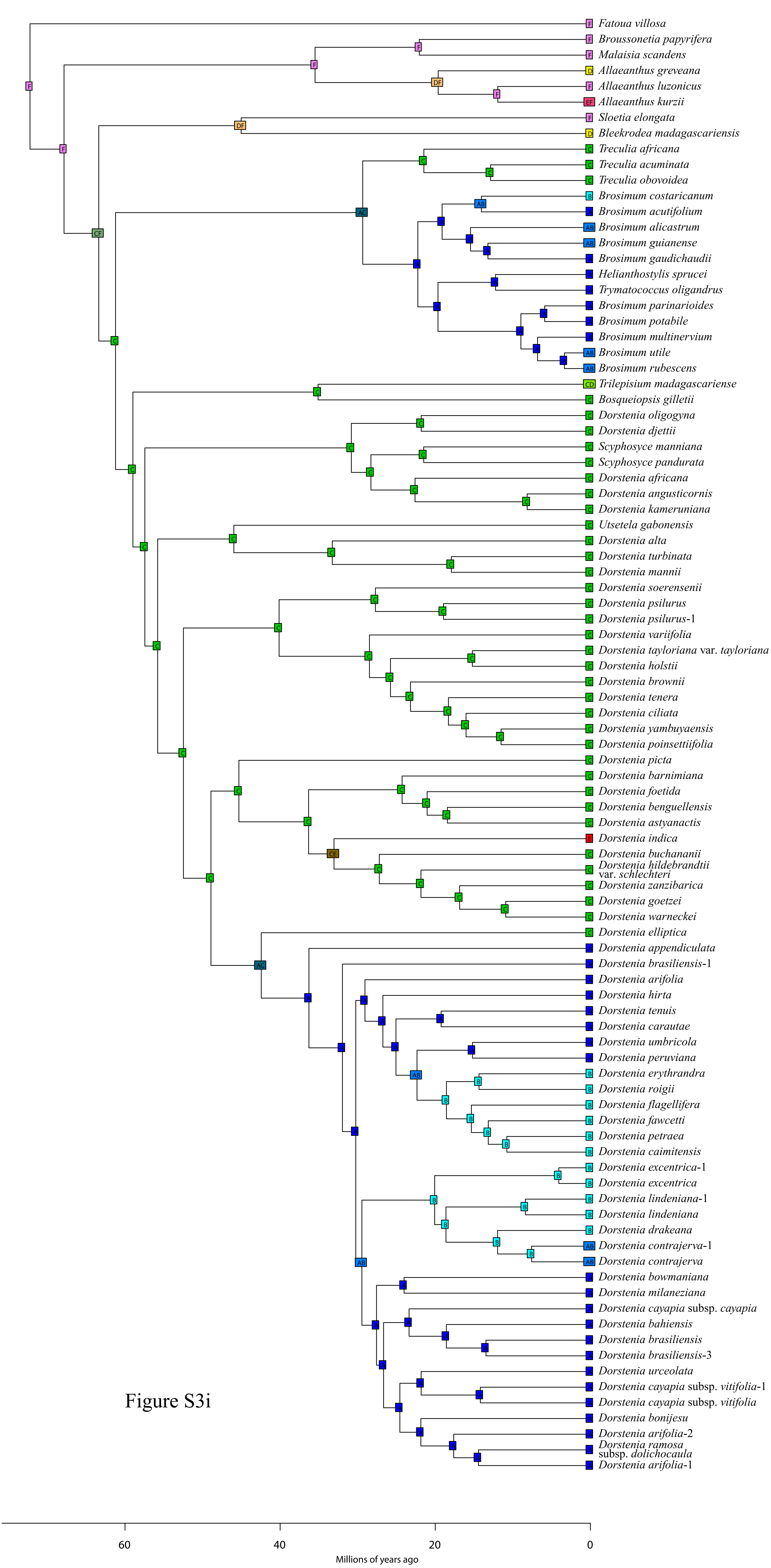


Figure S3i

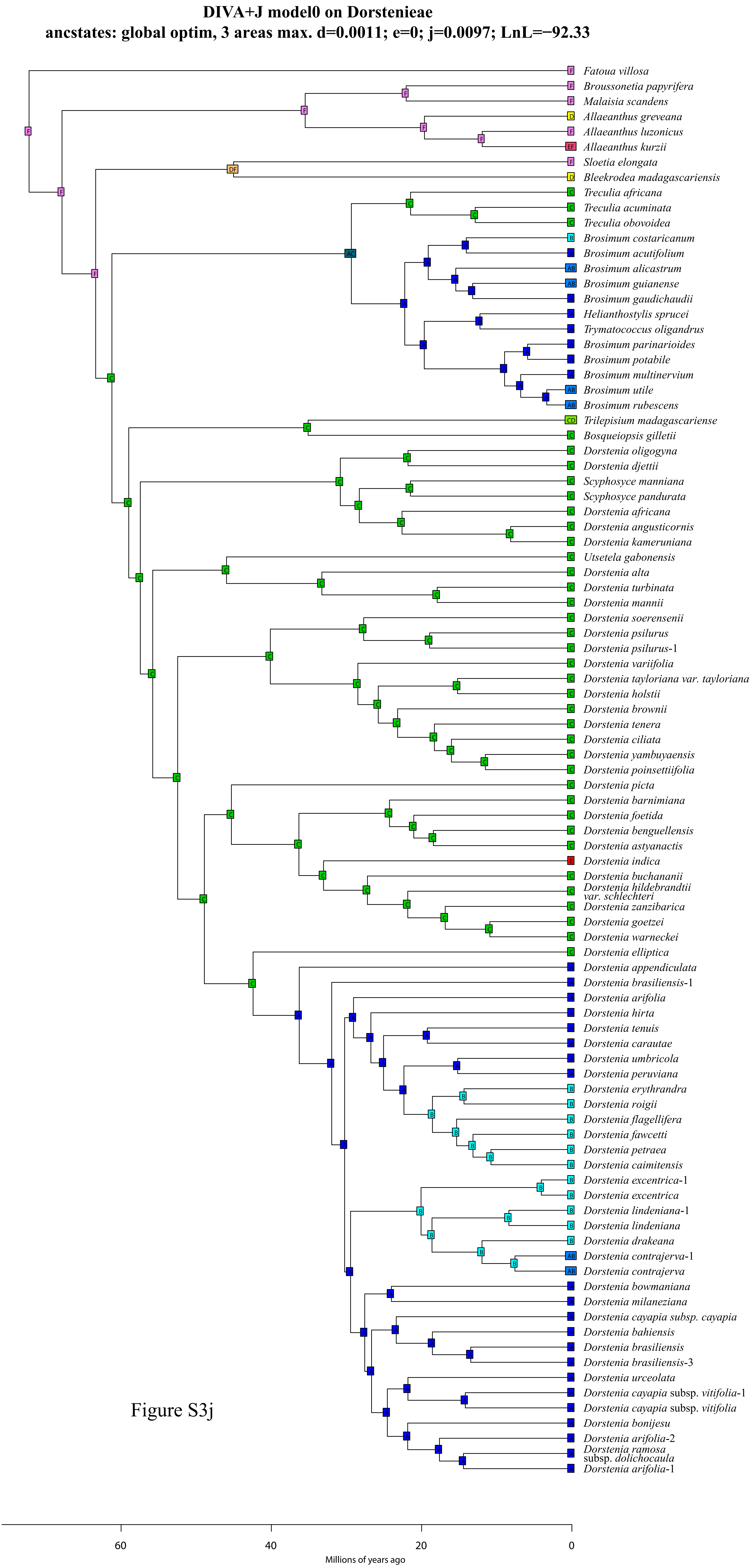
DIVA+J model0 on Dorstenieae
ancstates: global optim, 3 areas max. d=0.0011; e=0; j=0.0097; LnL=−92.33
Fatoua villosa
Broussonetia papyrifera
Malaisia scandens
Allaeanthus greveana
Allaeanthus luzonicus
Allaeanthus kurzii
Sloetia elongata
Bleekrodea madagascariensis
Treculia africana
Treculia acuminata
Treculia obovoidea
Brosimum costaricanum
Brosimum acutifolium
Brosimum alicastrum
Brosimum guianense
Brosimum gaudichaudii
Helianthostylis sprucei
Trymatococcus oligandrus
Brosimum parinarioides
Brosimum potabile
Brosimum multinervium
Brosimum utile
Brosimum rubescens
Trilepisium madagascariense
Bosqueiopsis gilletii
Dorstenia oligogyna
Dorstenia djettii
Scyphosyce manniana
Scyphosyce pandurata
Dorstenia africana
Dorstenia angusticornis
Dorstenia kameruniana
Utsetela gabonensis
Dorstenia alta
Dorstenia turbinata
Dorstenia mannii
Dorstenia soerensenii
Dorstenia psilurus
Dorstenia psilurus-1
Dorstenia variifolia
Dorstenia tayloriana var. tayloriana
Dorstenia holstii
Dorstenia brownii
Dorstenia tenera
Dorstenia ciliata
Dorstenia yambuyaensis
Dorstenia poinsettiifolia
Dorstenia picta
Dorstenia barnimiana
Dorstenia foetida
Dorstenia benguellensis
Dorstenia astyanactis
Dorstenia indica
Dorstenia buchananii
Dorstenia hildebrandtii var. schlechteri
Dorstenia zanzibarica
Dorstenia goetzei
Dorstenia warneckei
Dorstenia elliptica
Dorstenia appendiculata
Dorstenia brasiliensis-1
Dorstenia arifolia
Dorstenia hirta
Dorstenia tenuis
Dorstenia carautae
Dorstenia umbricola
Dorstenia peruviana
Dorstenia erythrandra
Dorstenia roigii
Dorstenia flagellifera
Dorstenia fawcetti
Dorstenia petraea
Dorstenia caimitensis
Dorstenia excentrica-1
Dorstenia excentrica
Dorstenia lindeniana-1
Dorstenia lindeniana
Dorstenia drakeana
Dorstenia contrajerva-1
Dorstenia contrajerva
Dorstenia bowmaniana
Dorstenia milaneziana
Dorstenia cayapia subsp. cayapia
Dorstenia bahiensis
Dorstenia brasiliensis
Dorstenia brasiliensis-3
Dorstenia urceolata
Dorstenia cayapia subsp. vitifolia-1
Dorstenia cayapia subsp. vitifolia
Dorstenia bonijesu
Dorstenia arifolia-2
Dorstenia ramosa subsp. dolichocaula
Dorstenia arifolia-1
60
40
20
0
Millions of years ago

Figure S3j

**DIVA model1 on Dorstenieae**

**ancstates: global optim, 3 areas max. d=0.0083; e=4e−04; j=0; LnL=−89.10**

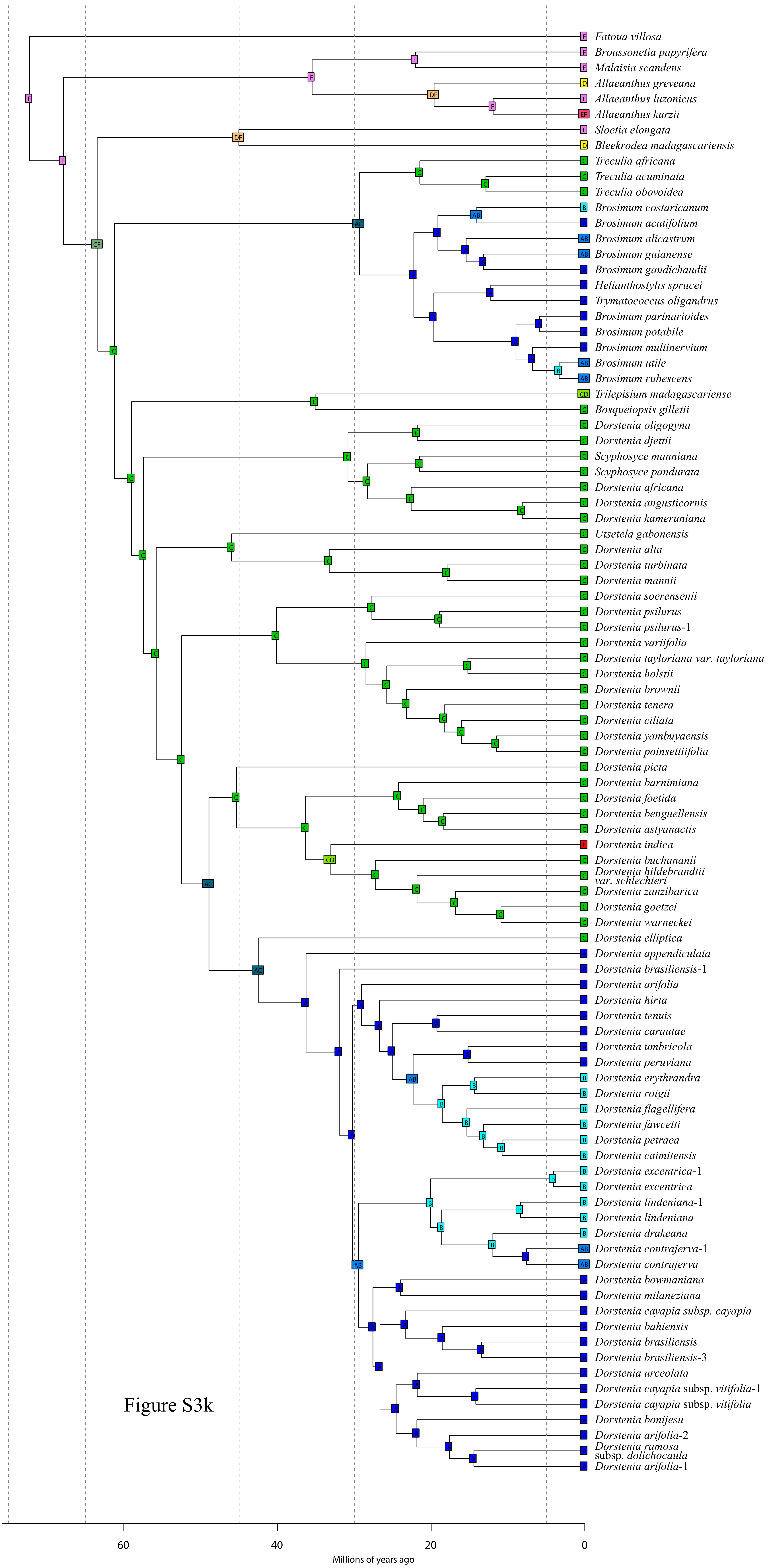


Figure S3k

DIVA+J model1 on Dorstenieae
ancstates: global optim, 3 areas max. d=0.0054; e=0; j=0.0485; LnL=−83.43

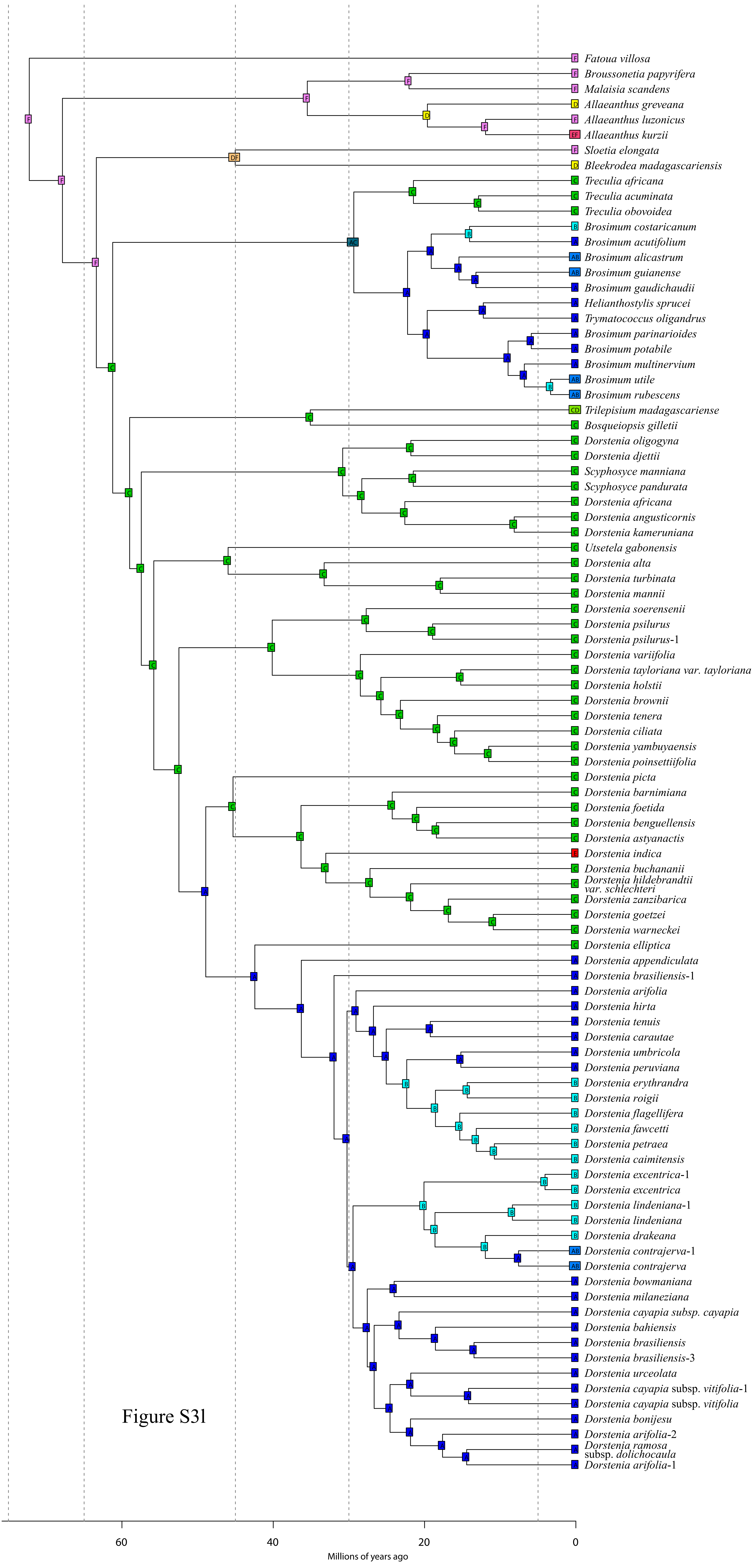


Figure S3l
